# Synchronizing speciation, extinction, and dispersal to island paleodynamics through Bayesian phylogenetics in a Hawaiian plant radiation

**DOI:** 10.64898/2025.12.16.694722

**Authors:** Isaac H. Lichter-Marck, Sarah K. Swiston, Fábio K. Mendes, Michael R. May, Suman Neupane, Bruce G. Baldwin, Kenneth R. Wood, Nina Rønsted, Warren L. Wagner, Felipe Zapata, Michael J. Landis

## Abstract

Islands are natural laboratories for studying dispersal, speciation, and extinction, yet the historical dynamics of many insular radiations remain poorly understood because of the difficulty aligning the tempo and mode of diversification with paleogeography during phylogenetic inference. We introduce a phylogenetic modeling framework, named TimeFIG, that combines data about species ranges and genetic variation with paleogeographic information to simultaneously infer divergence times, ancestral ranges, and historical diversification rates, without relying on fossils to time-calibrate the phylogeny. Applying TimeFIG to the spectacular radiation of Hawaiian *Kadua* (Rubiaceae) points to colonization of either modern Kaua‘i or ancient now-eroded islands, with island isolation as the strongest correlate of dispersal, and with a tendency for net diversification to be highest on “middle-aged” islands. Modeling diversification within an explicit spatiotemporal context, while accounting for historical uncertainty regarding the timing and placement of phylogenetic and biogeographic events, enables the rigorous testing of foundational hypotheses in island biology and evolutionary theory for clades beyond *Kadua*.

## INTRODUCTION

At more than 3,500 km from the closest continent, the Hawaiian archipelago is among the most isolated landmasses on Earth and yet, against all odds, it hosts a diverse, endemic biota^1^. Long-distance dispersal across the Pacific Ocean, insular diversification, and dynamic cycles of island growth and subsidence have populated the archipelago with a multitude of unique lineages^2,3^. These properties have made Hawai‘i instrumental for developing and testing classic evolutionary and ecological theories regarding the cascading effects of eco-geological change on the origin, evolution, and extinction of species^4–6^, such as the equilibrium theory of island biogeography^7^, the progression rule of island biogeography^3^, and the general dynamic theory of oceanic island biogeography^8^.

To evaluate support for the above theories, which concern the production of biodiversity patterns in space over time, biologists often employ phylogenetic methods to reconstruct the historical biogeography and evolution in clades of living species^9–11^. Unfortunately, phylogenetic studies of island taxa remain constrained by two long-standing challenges^12^: island radiations often lack fossils useful for establishing the geological timing of island colonization and diversification, and there are limited methods to jointly infer phylogenetic and biogeographic histories within a dynamic paleogeographic context. These challenges make it difficult to infer the tempo and mode of island lineage diversification, thus weakening our understanding for how geography and environment shape island radiations through time.

Studying the detailed historical biogeography for island clades generally relies on one of three popular strategies: assume the clade is younger than the oldest island it occupies^13^, at the risk of introducing circularity into the study^14^; use fossil evidence associated with deeply ancestral taxa to pre-estimate the age of the island clade^15,16^, while ignoring the influence of island paleogeography on lineage divergence^17^; or use non-phylogenetic methods^18^, which do not tap the potential of phylogeny for testing biogeographic hypotheses^19^. While much can be learned using any of these three strategies, an alternative approach might profit from jointly analyzing all information that unifies the phylogenetic, biogeographic, and paleogeographic histories of the study system^20^.

To this end, we design a time-heterogeneous phylogenetic model of biogeographic diversification (TimeFIG), which uses paleogeographic information to simultaneously reconstruct phylogenetic relationships, divergence times, ancestral ranges, and biogeographic rates of dispersal, speciation, and extinction. We then apply TimeFIG to study the diversification of Hawaiian *Kadua* (Rubiaceae)^21^. Comprising approximately 25 extant taxa distributed across all modern Hawaiian island complexes (i.e., Kaua‘i, O‘ahu, Maui Nui, Hawai‘i), *Kadua* is ideal for investigating the timing of island colonization and diversification within a paleodynamic landscape^22^. Previous phylogenetic studies inferred a contentious arrival date for *Kadua* into the archipelago between 3–13 Ma, potentially supporting recent colonization coincident with the emergence of Kaua‘i, or older colonization of the now-eroded Northwestern islands^23^. Which island characteristics fueled the extensive radiation of *Kadua* throughout the archipelago and across new adaptive zones, while producing remarkable disparity in life history, reproductive mode, dispersal strategy, and habitat occupancy^21,23^ remain open questions, such is the case for many insular plant lineages.

We used TimeFIG to test four specific, mutually compatible biogeographic hypotheses: smaller islands have higher extinction rates (H1: the size-extinction relationship^7^); dispersal is negatively associated with island isolation (H2: the distance-dispersal relationship^7,24^); dispersal is higher from older to younger islands (H3: the progression rule of island biogeography^3^); and net diversification varies temporally with the growth and the erosion and subsidence of islands (H4: dynamic island biogeography^8^).

## RESULTS

### Time-calibrating island radiations with paleogeography

TimeFIG allows time-varying regional features (e.g., size, distance) to shape biogeographic rates of speciation, extinction, and dispersal during phylogenetic inference by using feature effect parameters “(Figure 1)”. For instance, larger islands may experience higher within-region speciation rates than smaller islands when a feature effect parameter governing this relationship is positive, 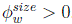. Analogously, larger islands would have slower rates if 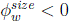, and island size would have no impact if 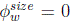. Such relationships are maintained even as regional features change over time, e.g., speciation accelerates as islands grow when 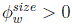. More generally, TimeFIG allows measurements associated with each regional feature, *f*, to have positive, negative, or no effect on rates of <u>d</u>ispersal, <u>e</u>xtinction, <u>w</u>ithin-region speciation, or <u>b</u>etween-region speciation, indexed with *p* ∈ {*d, e, w, b*}, by using feature effect parameters 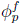 to model relationships with quantitative features (e.g., size, distance) and 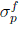 for categorical features (e.g., big vs. small).

**Figure 1.**
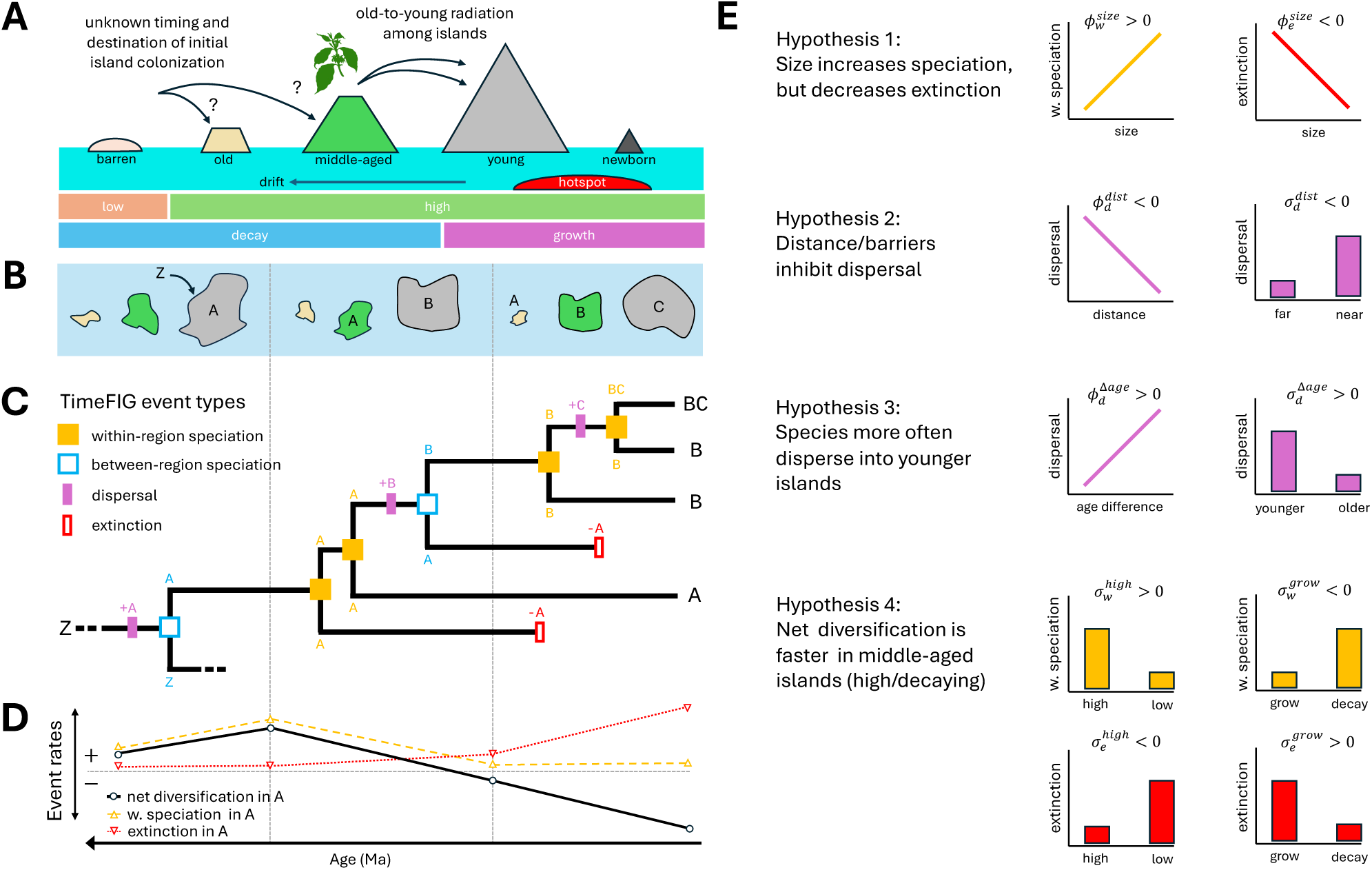
Hypothesized TimeFIG behavior for volcanic island biogeography. (A) Volcanic islands transition through predictable phases of growth and decay after origination. The exact timing and location of island colonization events is not known. (B) Paleogeographic system showing how volcanic islands grow while over the hotspot, but subside and decay as they drift away from it over time. Focal islands — A, B, C — are initially colonized from the external region, Z. (C) TimeFIG models biogeographic radiation within the paleogeographic system using four event types: within-region speciation (filled, gold square), between-region speciation (open, blue square) dispersal (filled, purple rectangle), and extinction (open, red rectangle). Example scenario shows an island radiation unfolding over three time periods. First, island A is colonized from external region Z. Next, the lineage diversifies in A and later colonizes the newly formed island B. Lastly, most species in A go extinct as the island grows barren, while species in B diversify and colonize the newly formed island C. (D) Example TimeFIG rates of net diversification (black), within-region speciation (gold), and extinction (red) for region A that are compatible with scenarios shown in Panels B and C. (E) Expected relationships between TimeFIG feature effect parameters (*ϕ*^(*k*)^ or *σ*^(*k*)^) and regional features for the four focal hypotheses of this study. Briefly, positive values for feature effect parameters indicate event rates and regional features are positively correlated, whereas negative values indicate a negative correlation. See main text for more details. Image of plant from https://phylopic.org.

Importantly, TimeFIG assumes effects of regional features on biogeographic rates are unknown, and are estimated from the phylogenetic relationships, divergence times, the presences and absences of each species in each region, and paleogeographic information. For instance, many young species on a large island might indicate the positive effect of region size on within-region speciation (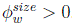). Conversely, few species on old islands is consistent with a negative effect of region age on extinction (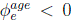), but also with a negative effect of age-difference between islands on dispersal (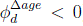). Support for all feature effects are estimated jointly from the data, which collectively induce rate variation across all regions, time periods, and biogeographical processes, while using relatively few model parameters. For our Hawaiian system, which involves 7 time periods, 7 regions, and 16 regional features, TimeFIG uses 20 parameters (4 base rates, 16 feature effects) to model biogeographical rate variation, whereas a standard geographic state-dependent speciation-extinction (GeoSSE^16^) model with time-varying rates requires hundreds of parameters to attain equivalent levels of variation.

TimeFIG assigns different probabilities to phylogenetic and biogeographic patterns based on how well they fit to paleogeographic data, producing molecular phylogenies that are realistically scaled to geological time^25^ “(Figure S1)”. When analyzing simulated Hawaiian radiation datasets with young colonization times (1 to 6 Ma), TimeFIG accurately estimated the true age of the island radiation in 82% of cases, compared to 1% of cases for analyses under an uncalibrated birth-death model (“Figure S3”, top). For older colonization times (6 to 20 Ma), TimeFIG correctly identified the true clade age in 83% of cases, compared to 40% of cases for those uncalibrated analyses with older, true ages that, by chance, matched the model’s uninformed expectations (“Figure S3”, bottom).

Analyses of simulated Hawaiian radiations also revealed that TimeFIG can reliably detect many, but not all, of the feature effects related to our four biogeographic hypotheses (listed in “Figure 1E” and “Table S4”). Highest posterior density (HPD) intervals more often contained true effect values and/or excluded the incorrect sign for a hypothesis in datasets where the targeted effect was truly present (“Figure S4”) versus those where it was absent (“Figure S5”, “Table S7”), except for five parameters that were generally difficult to estimate (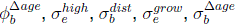). Assuming Bayes factors (BFs) with values *>*3.2 indicate ‘substantial’ significance^26^ “(Figure S6; Table S8)”, six effects (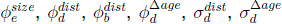) were correctly detected in *>*50% of cases, six effects (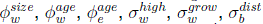) were detected in 15-50% of cases, and four effects were detected in *<*15% of cases (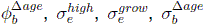). Encouragingly, false discovery rates (FDRs) were low (*<*15%) for all effects, excluding five effects with moderate FDRs (15-30%) (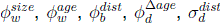), and four effects with unknown FDRs (too few true/false discoveries; 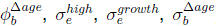). Using larger Bayes factors to determine significance drove most FDRs to 0% “(Figure S6; Table S8)”.

### Hawaiian *Kadua* radiation: surviving only among modern “high” islands

Using 344 nuclear loci, we inferred the phylogenetic tree topology of *Kadua* by sampling 63 individuals and 5 outgroup taxa distributed across all modern Hawaiian “high” island complexes, French Polynesia, and continental Asia (“Figure 2A-E”), capturing 23 of 25 currently recognized Hawaiian species (“Figure 2F-G”). Fourteen species occur on the oldest modern “high” island, Kaua‘i, with six being endemic. In contrast, only seven species occur on the youngest “high” island, Hawai‘i, all of which are widespread across multiple islands, suggesting recent range expansions.

**Figure 2.**
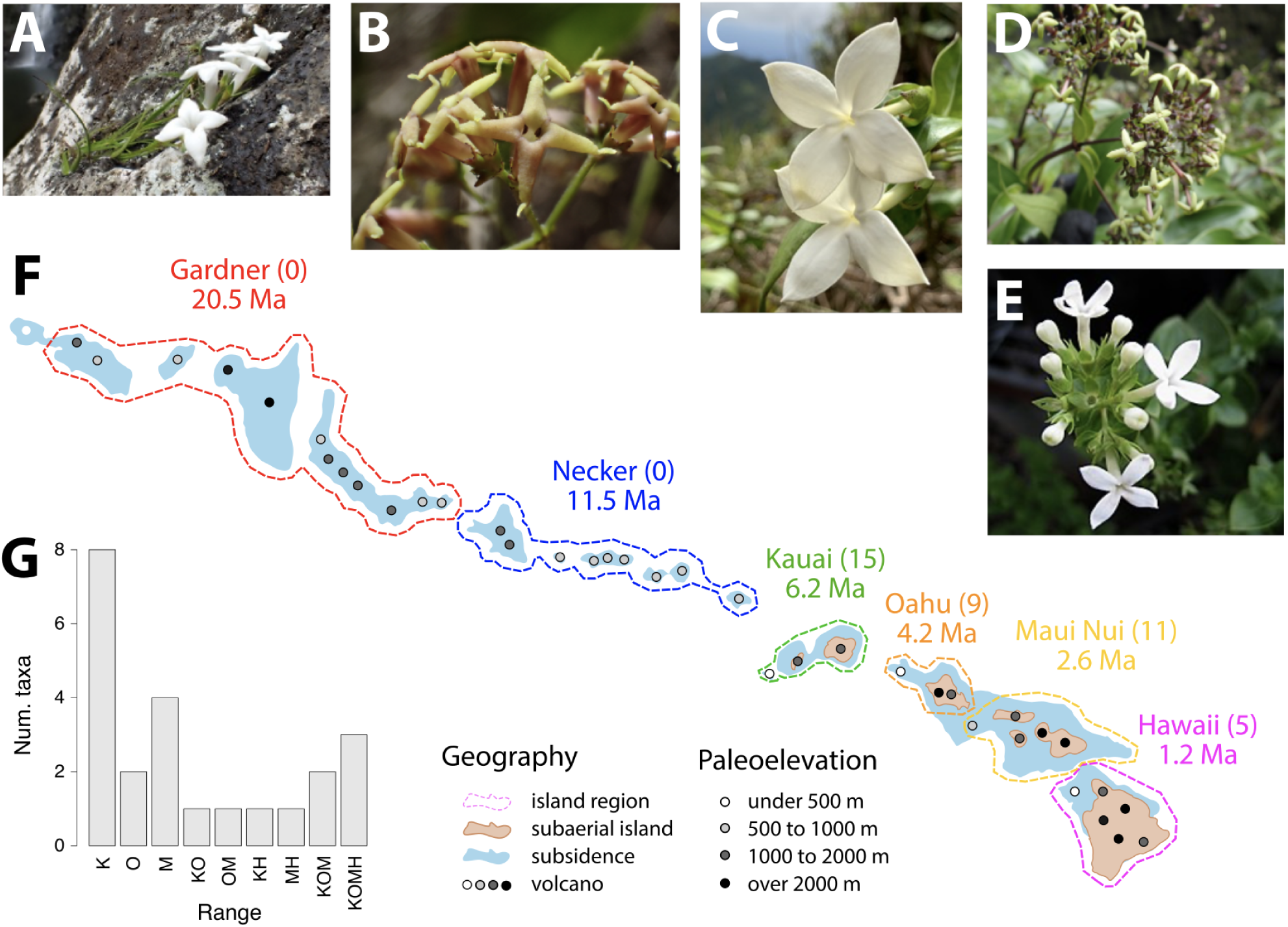
Extant Hawaiian *Kadua* only occupy the modern “high” islands. (A-E) Representatives of the Hawaiian *Kadua* radiation: (A) *K. cookiana* (B) *K. knudsenii* (C) *K. fluviatilis* (D) *K. centranthoides* (E) *K. parvula*. (F) The radiation contains 25 taxa distributed throughout the Hawaiian archipelago. Islands within the archipelago follow a “conveyor belt” pattern, with islands originating in the southeast, then drifting to the northwest. Volcanism and uplift followed by subsidence and erosion generally caused island sizes to increase then decrease, portrayed by the ancestral (blue) and current (brown) island sizes and the maximum elevation of ancestral volcanoes (shaded circles). Islands are grouped into six regions (dotted lines), with each region appearing at the time of the oldest island in the complex. The numbers of taxa per region (in parentheses) show that native extant *Kadua* only inhabit the modern high islands (Hawai‘i, pink; Maui Nui, yellow; O‘ahu, orange; Kaua‘i and Niihau, green), with none found among the older northwestern islands (Necker complex, blue; Gardner complex, red). (G) Of the 25 taxa, 14 taxa occur in one island region, and 9 occur in multiple regions. Polynesian Islands and outgroup taxa (region Z) are not shown. Images of *Kadua* by K. Wood. Image for Hawaiian islands modified from Price et al.^27^ and Givnish et al.^13^.

Although no living *Kadua* species inhabit the nearly barren Northwestern islands predating Kaua‘i, Gardner and Necker islands once rivaled today’s modern “high” islands in terms of size and elevation (“Figure 2F”)^27^, plausibly providing suitable habitats for ancestral species. Importantly, the absence of extant *Kadua* from Necker and Gardner today is itself a source of information rather than missing data: each taxon is explicitly scored as absent from these regions, so the analysis treats present-day absences as evidence that any ancestral lineages, if once present there, subsequently either went extinct or dispersed elsewhere. This evidence of true absence – rather than lack of observation – helps TimeFIG weighs historical scenarios in which *Kadua* might have once occupied the now-eroded islands.

Our results support a monophyletic *Kadua* that colonized the Hawaiian archipelago and underwent extensive insular diversification (“Figure 3A”). Three French Polynesian species nested within *Kadua* indicate an out-of-Hawai‘i dispersal event (“Figures S7-S8”). The *Kadua* relationships we recovered reinforce patterns found in previous molecular work^23^, and align with taxonomic hypotheses proposed by Terrell et al.^21^ based on morphology. Concatenation and coalescent-based analyses produced similar phylogenetic relationships, except within the highly nested clade *Kadua* sect. *Wiegmannia* (“Figure S9”). Our results further support the monophyly of most *Kadua* species, except the widespread *K. cordata* species complex, within which we recover four independent lineages, including one undescribed cliff-dwelling taxon requiring further study.

**Figure 3.**
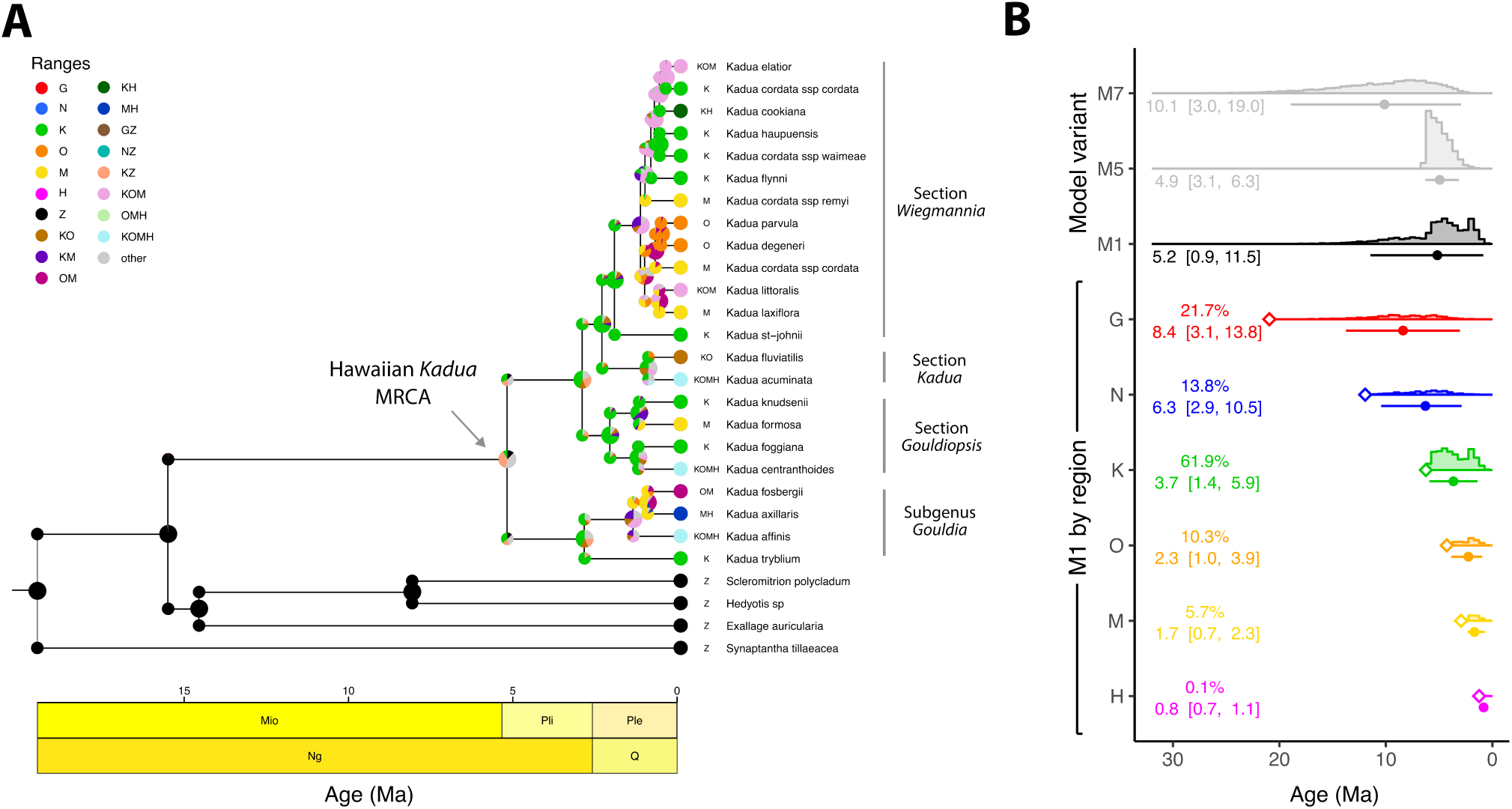
Phylogenetic and biogeographic reconstruction for *Kadua* lineages in the Hawaiian archipelago. (A) Ancestral range estimates show that the most recent common ancestor of all surviving Hawaiian *Kadua* (gray arrow) most likely occupied Kaua‘i [K] and/or the external region [Z]. (B) Posterior age estimates for the Hawaiian *Kadua* MRCA when using no time-calibration strategy (M7), when using a biogeographic node age constraint (M5), or when using the full TimeFIG model (M1). Circles represent posterior means and bars represent 95% highest posterior densities (HPD95). Age estimates under TimeFIG (M1) are further separated by island regions in the ancestral range of the *Kadua* MRCA. Diamonds indicate when each island region first appeared. Percents report the marginal probability the MRCA occupied the island region; species ranges can include multiple regions, so these percents do not sum to 100%. Numbers in brackets report the HPD95 intervals.

### Co-estimating when and where *Kadua* arrived in Hawaii

We used TimeFIG to date when and where the Hawaiian *Kadua* radiation began (“Figures 3 and S10”, “Table S9”). The posterior mean age of the most recent common ancestor (MRCA) of extant *Kadua* was ∼5.2 Ma, which is younger than any modern island (∼6.2 Ma). Mean estimates of MRCA ages of the four major subclades were approximately 0.88, 2.0, 2.8, and 2.9 Ma, all younger than O‘ahu (∼4.1 Ma), the second-oldest modern island complex. While all four subclades almost certainly originated among the modern “high” Hawaiian islands, our results suggest that the *Kadua* MRCA might have first diversified among the older Necker (*p* = 0.14) and/or Gardner (*p* = 0.22) regions (“Figure 3B”).

Age estimates for the MRCA of *Kadua* differ substantially when inferred using a process-calibrated TimeFIG model (M1), a node-calibrated birth-death model (M5), or a molecular clock with an uncalibrated birth-death model (M7; “Figure 3B”; models M1–M8 shown in “Figure S11”). The uncalibrated analysis contains little or no information to estimate the true age of the island clade and consequently exhibit the highest uncertainty (posterior mean and HPD95 of 10.1 [2.9, 18.9] Ma). The node-calibrated analysis assigns zero probability to *Kadua* being older than Kaua‘i, effectively truncating the upper bound of the M7 estimates (4.9 [3.1, 6.3] Ma), but would introduce circularity into downstream biogeographical analyses^28^. In contrast, the process-calibrated TimeFIG analysis estimates younger ages while permitting older, pre-Kaua‘i origination times (5.2 [0.9, 11.5] Ma), while simultaneously estimating ancestral ranges and rate dynamics.

Multimodal clade age estimates under TimeFIG reflect inherent uncertainty about when and where *Kadua* first arrived and radiated within the archipelago (“Figure 3B”). Younger clade ages are chiefly associated with Kaua‘i ancestry (3.7 [1.4, 5.9] Ma, marginal probability of regional occupancy is *p* = 0.62) and secondarily with O‘ahu (2.3 [1.0, 3.9] Ma, *p* = 0.10) or Maui Nui (1.7 [0.7, 2.3], *p* = 0.06), whereas older clade ages are associated with Gardner (8.4 [3.1, 13.8] Ma, *p* = 0.22) and Necker (6.3 [2.9, 10.5] Ma, *p* = 0.14) ancestry. (Note, because species can occupy multiple islands, these marginal probabilities across islands need not sum to 1). Histories where the MRCA of *Kadua* includes region (Z) represent scenarios where island diversification began immediately following colonization, or with multiple colonizations of the archipelago (e.g., from Hawaiian islands older than Gardner, other Pacific islands, or a continental mainland).

### Island isolation and age shape biogeographic rates

Inference of 16 relationships between biogeographic rates with quantitative (“Figure 4”, left) and categorical (“Figure 4”, right) island paleofeatures revealed three characteristics that strongly influenced *Kadua* dispersal and diversification, and four features that displayed weaker influence (“Figure 4”, middle; “Tables S10-S11”). Our estimates weakly support the hypothesized association between smaller islands having increased extinction rates^7^ (posterior mean and HPD80 of 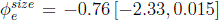; BF 4.0), though no significant relationship was detected between island sizes and speciation rates (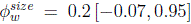; BF 1.5). We find strong support for dispersal rates decreasing with geographical distance (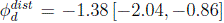; BF 97.0)^7^ and for less transoceanic dispersal entering and leaving the archipelago (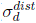 = −2.00 [−3.04, −1.06]; BF 147.0)^24^. Support for the progression rule^3^ was weak, but requires subtle interpretation. Dispersal favored movement from older into younger islands as age differences increased (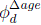 = 0.50 [0.00, 1.14]; BF 3.7), but was partly counterbalanced by slightly higher back-dispersal rates from younger into older islands, regardless of the exact age difference (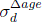 = −0.19 [−1.05, 0.31]; BF 0.5). Our findings also aligned with predictions that diversification accelerates during the early stages of island growth and tapers off during island decay^8^. Within-region speciation rates were elevated for islands in the high-island phase (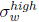 = 0.90 [0.00, 1.76]; BF 7.3), but partly reduced during the “net growth” phase (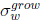 = −0.29 [−1.10, 0.00]; BF 2.2), tending to produce the highest rates for middle-aged islands that were partly eroded and subsident, but still high. No relationships were detected between extinction and high-island (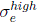 = −0.05 [−0.89, 0.61], BF 1.0) or net-growth status (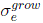 = 0.04 [−0.78, 0.70], BF 1.0). When conditioning on *Kadua* being older than Kaua‘i, the negative impact of island size on extinction strengthens considerably (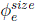 = −0.76 [−2.92, 0.00]; BF 22.0) compared to our ‘any-age’ estimate, and lacks significance when conditioning on *Kadua* being younger than Kaua‘i (−0.49[−1.93, 0.40]; BF 2.7). Based on our simulations, the false discovery rates of all signed effects at their respective significance levels is low (FDR≤5.0%), except for 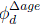 (FDR=26.1%; “Table S11”). In summary, Bayes factors indicate substantial support (BF*>*3.2) for at least one parameter pertaining to each of the four hypotheses (“Figure 1E”, “Table S4”): 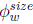 for H1, 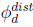 and 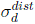 for H2, 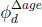 for H3, and 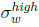 for H4.

**Figure 4.**
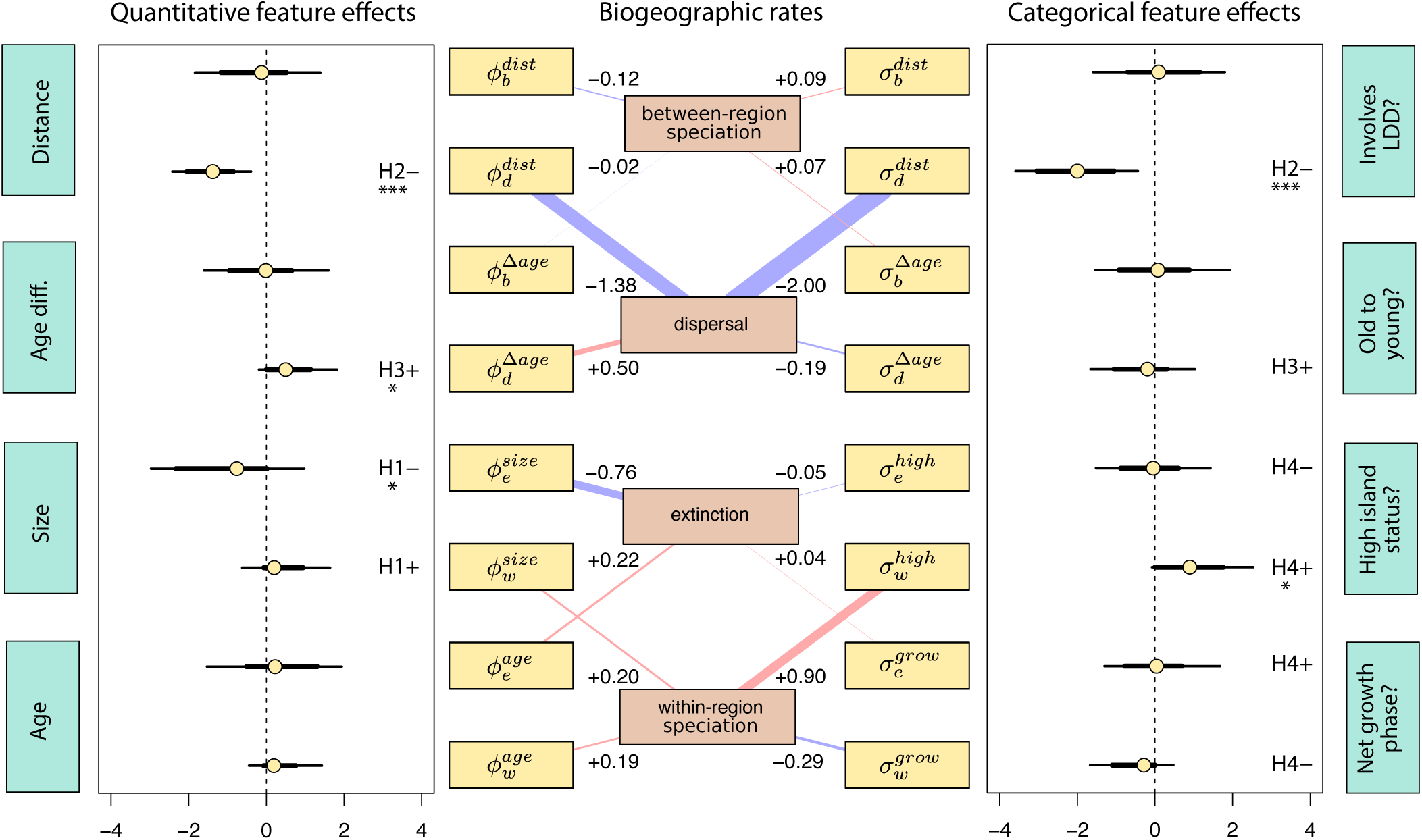
Effects of regional features on biogeographic rates. Estimates account for all phylogenetic and biogeographic histories supported by the model. Marginal posterior densities show the sign and scale of relationships between regional features and biogeographic rates through 8 quantitative (left) and 8 categorical (right) feature effect parameters. For the left and right flanking plots, points represent posterior means, thin lines represent 95% HPD intervals, and thick lines represent 80% HPD intervals. Parameters are annotated with predicted positive (+) and negative (–) relationships under our four biogeographic hypotheses (H1, H2, H3, H4; see “(Figure 1)”). Asterisks indicate substantial (*; *>*3.2), strong (**; *>*10), or very strong (***; *>*32) Bayes factor support for each signed hypothesis. For the middle plot, edges in the network summarize posterior mean strength (edge width) and sign (blue: negative, red: positive) of relationships between regional features (cyan labels) and biogeographic event rates (brown labels) through their associated feature effect parameters (gold labels).

### Net diversification tends to be highest on middle-aged islands

The collective effect of all paleofeatures likely caused speciation and extinction rates for Hawaiian *Kadua* to rise and fall as the islands experienced uplift, erosion, and subsidence (“Figure 5”, “Figure S12”, “Table S12”). The trajectory of the Gardner island complex best illustrates how net diversification rates – the difference between speciation and extinction rates in region *i* during time interval *t*, given as *r_w_*(*i, t*) − *r_e_*(*i, t*) – vary over time. Before island birth, we assume net diversification rates are zero (*r_v_*(*G,* 7) = 0.0), even though it would be negative for a submerged island. Upon emergence and throughout the early growth phase, net diversification rates for Gardner begin lower but positive (posterior median and HPD50 for *r_v_*(*G,* 6) = 0.24 [0.05, 0.45]), rising to its maximum when this then-high island first entered its net decay phase (*r_v_*(*G,* 5) = 0.28 [−0.10, 0.78]). As large high islands age, becoming small low islands, diversification rates decline (*r_v_*(*G,* 4) = 0.16 [−0.48, 1.00]), continuing into the final stages of island maturity, when net diversification rates tend to be negative and the islands are flat, ecologically simple, and nearly barren (*r_v_*(*G,* 2) = −0.19 [−3.05, 0.75]). Critically, these dynamics are completely absent from the induced prior distribution of the model, meaning they are inferred from the data rather than forced by TimeFIG (“Figure S13”).

**Figure 5.**
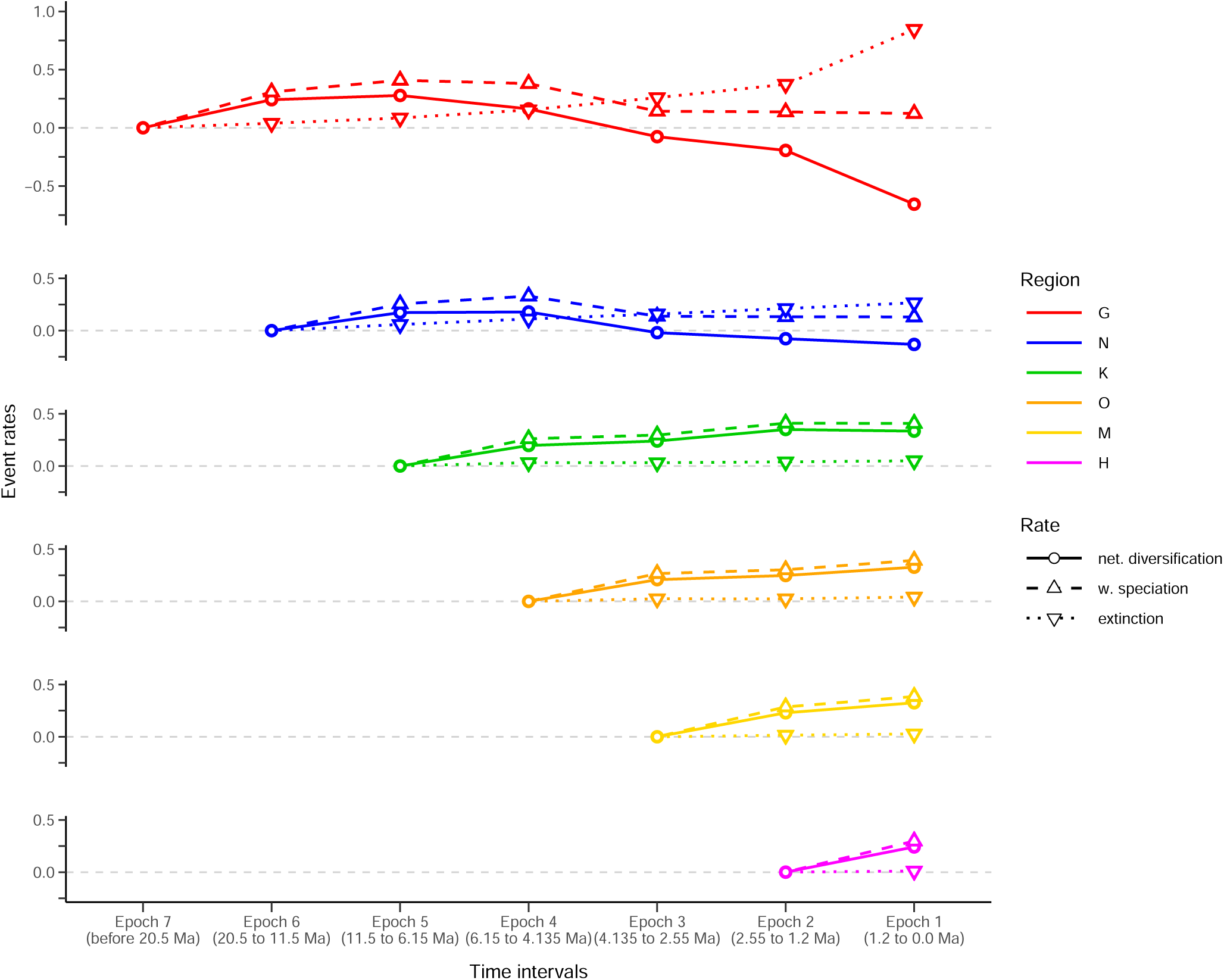
Regional rates of biogeographic processes over time. Estimates are averaged over all phylogenetic and biogeographic histories supported by the model. Posterior median net diversification rates (*r_w_*(*i, t*) − *r_e_*(*i, t*); circles), within-region speciation rates (*r_w_*(*i, t*); upward triangles), and extinction rates (*r_e_*(*i, t*); downward triangles) in region *i* during epoch *t* from past (left) to present (right) at different stages of Hawaiian paleogeographic history. Rate variation among time intervals and regions is determined by feature effect parameter values interacting with paleogeographic changes in underlying regional features. The net diversification rate is assumed to be zero for submerged islands. Note that all panels share the same y-axis scale, except for Gardner (top). The external region, Z, is not shown.

In general, regional net diversification rates tend to rise then fall as islands mature, advancing from being young to middle-aged to old (“Figure 6”). Island birth and initial growth is linked to increased net diversification (mean change of +0.17 rate-units), with sustained increases throughout middle-age as islands mature (+0.05 rate-units), before rates decline in old age (−0.18 rate-units), eventually turning negative. Contemporary island regions provide a snapshot of the main stages of island life. The youngest island region of Hawai‘i currently has the lowest net diversification rate of all modern islands, but possesses the greatest potential for increased diversification in the near future. O‘ahu and Maui Nui, now middle-aged, might sustain positive net diversification rates for another 1-2 million years. Kaua‘i, the oldest modern island complex, is exiting its middle-aged period, with a high net diversification rate comparable to other modern islands that is now in subtle decline. Lastly, the ancient Gardner and Necker complexes, which contain no extant *Kadua* species today, now experience net species loss, with increasingly negative net diversification rates. If *Kadua* first colonized Gardner or Necker, its descendants would have needed to disperse into a younger island to survive.

**Figure 6.**
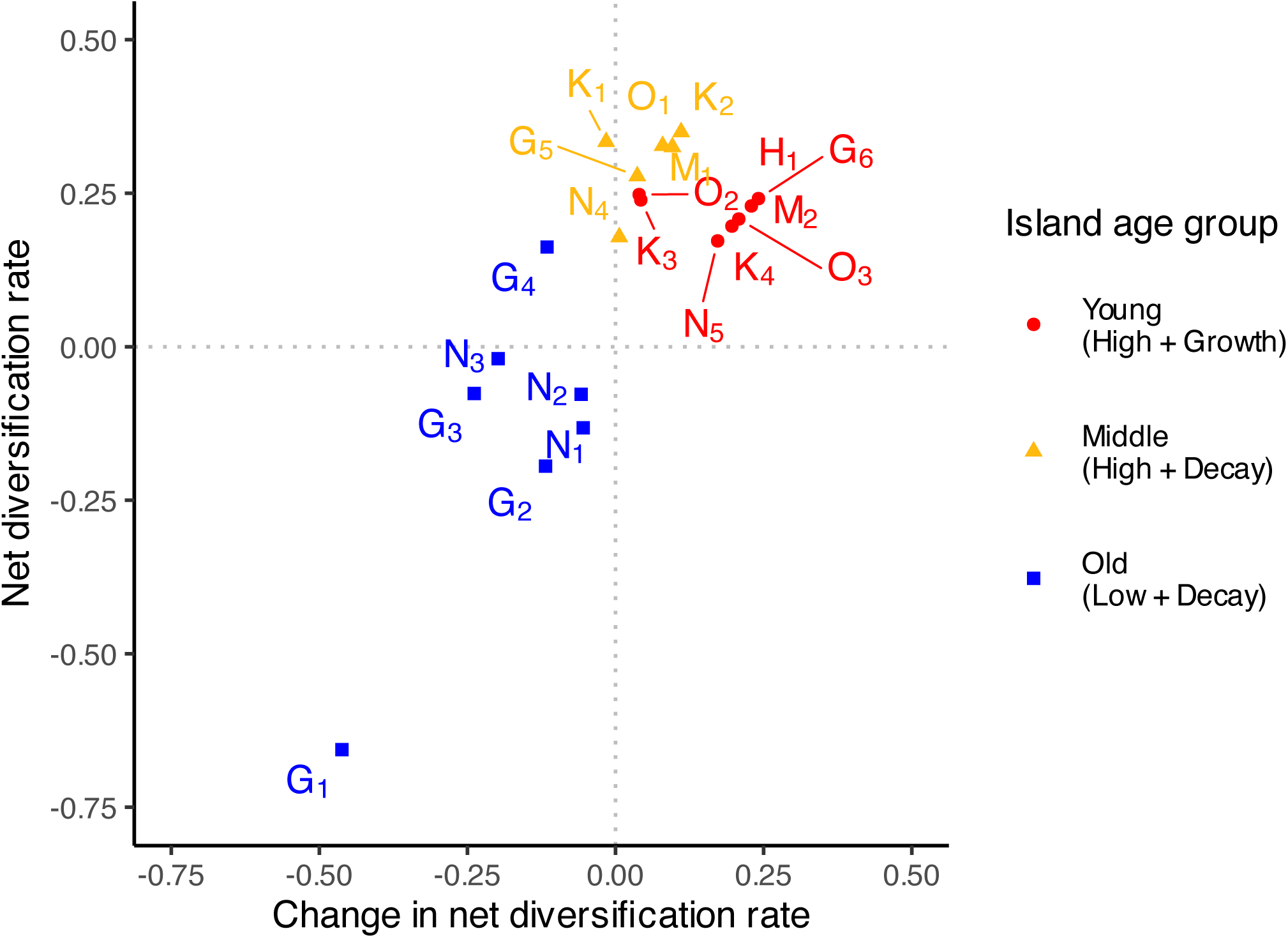
Middle-aged islands tend to have higher net diversification rates. Points represent the net diversification rates in region *i* during epoch *t* (*r_v_*(*i, t*) = *r_w_*(*i, t*) − *r_e_*(*i, t*)) versus the change in net diversification rates for region *i* between epochs (*r_v_*(*i, t*) − *r_v_*(*i, t* + 1)), where epoch *t* + 1 precedes epoch *t* (see “Figure 5”) The net diversification rate is assumed to be zero for submerged islands. Island at each epoch are classified as young (high + uplift; red), middle-aged (high + decay; gold), or old (low + decay; blue), and are labeled by letter and epoch (subscript). Young and middle-aged islands (except Kaua‘i at the present) have increasing net diversification rates, while all old islands have decreasing rates.

## DISCUSSION

### The colonization and radiation of Hawaiian *Kadua*

Using sequence target capture, we estimated a resolved phylogeny of *Kadua* (“Figures S7-S9”), illustrating the power of genomics for inferring relationships among island plant lineages (see also:^9,10,29–31^). Hawaiian *Kadua* was recovered as paraphyletic with respect to species from French Polynesia and the Austral Islands^23^, suggesting that *Kadua* first colonized the Hawaiian archipelago, presumably over the Pacific Ocean from Indo-Malaysia, then subsequently colonized the Polynesian islands^2^, as reported in other Hawaiian plant groups^32–34^. We found strong support for four clades of Hawaiian *Kadua* (“Figure 3”) corresponding to earlier proposed taxonomic sections based on fruit type, seed shape, and seed coat surface micro-anatomy^21^. Species richness in *Kadua* arose through the parallel diversification of these clades, each initially dispersing from older to progressively younger islands and speciating *in situ*, with numerous species becoming widespread, and some descendants back-dispersing into older islands (“Figure 3”). This nested insular radiation pattern resembles that of other Hawaiian and non-Hawaiian adaptive radiations, including lobelioids^10^, silverswords^35^, *Tetragnatha* spiders^36^, *Anolis* lizards^37^, and Andean *Espeletia* ^38^.

Using TimeFIG, we inferred probable scenarios for the arrival and expansion of *Kadua* in Hawai‘i (“Figure 3B”). The most probable scenario places *Kadua* in Kaua‘i during 1.4–5.9 Ma, with fair odds of it instead first colonizing the older island complexes, Necker and Gardner, during 2.9–13.8 Ma. Although the older colonization scenario is less probable than the Kaua‘i-first scenario, it is most compatible with previous fossil-based estimates informed by backbone lineages in tribe Spermacoceae of Rubiaceae^23^. Similarly, numerous Hawaiian insect clades are estimated as older than the modern islands^11,39,40^, yet evidence for ancient plant colonization remains scarce, with lobelioids being a notable exception^10,13^. Although our study cannot decisively conclude whether *Kadua* is younger or older than Kaua‘i, TimeFIG explicitly integrated over this historical uncertainty when testing our biogeographic hypotheses.

Our results show that island growth, erosion, and subsidence, which altered island sizes and topographies over time, plausibly impacted historical rates of dispersal and lineage diversification in Hawaiian *Kadua* (“Figure 4”). Island size, predicted to strongly influence extinction, exhibited a significant but weaker effect than anticipated^7,24^. This is surprising given the eight-fold size disparity between the youngest (Hawai‘i) and oldest (Kaua‘i) modern islands. Despite its relatively small size, Kaua‘i remains an ongoing center of *Kadua* diversification, harboring at least five speciation events during the past two million years (Sect. *Wiegmannia* in “Figure 3A”). We expect that size effects on diversification may not be apparent until much more extensive erosion and subsidence occur^8^. Indeed, islands that have almost entirely eroded away, such as the Gardner and Necker complexes, have the fastest extinction rates and contain no extant *Kadua*, likely reflecting local climate and ecosystem collapse that imperils resident species (“Figure 6”).

Distances between regions and isolating geographic barriers exhibited the strongest effect on dispersal among all features we examined, aligning island biogeography theory^7,24^. This pattern is common in other island systems^35,41^ and likely reflects limited organismal dispersal capabilities across long distances. Consistent with the progression rule^3^, *Kadua* lineages also generally dispersed from older to younger islands, particularly as age differences increased. It appears many Hawaiian lineages follow this rule^3,35,42,43^, potentially due to greater ecological opportunities in younger islands. Despite the old-to-young dispersal trend in *Kadua* being significant, simulations suggest there is a modest chance is a false positive (Table S11) and the effect strength is minimal (“Figure 4”), likely dampened by successful back-dispersal to older islands, as resolved in other Hawaiian plant lineages^10,44^.

*Kadua* diversification and island ontogeny appear coupled, consistent with the general dynamic theory of oceanic island biogeography^8,18^ “(Figure 5)”. Net diversification rates climaxed when islands reached middle age, after developing complex topography and diverse habitats, but before significant erosion and subsidence led the islands to become low-elevation and nearly barren “(Figure 6)”. Because signal rests on a single small clade and feature effects with modest strengths, we regard these results as suggestive rather than definitive. The temporal dynamics suggest that ecological opportunity provided by mature high islands through increased habitat heterogeneity and environmental gradients could drive *Kadua* diversification. The evolutionary response of *Kadua* to such opportunities is reflected in its radiation across diverse habitats, ecological niches, and morphological forms. Whether elevated net diversification rates among middle-aged islands^8^ generalizes across other Hawaiian lineages remains an open question that is addressable using our phylogenetic approach. More broadly, this underscores the importance of incorporating regional features into diversification studies, particularly for oceanic archipelagos, where ecological opportunity and evolutionary potential shift with the changing landscape.

Although our analyses affirmed several predicted relationships between paleofeatures and biogeographic rates, these relationships may not be directly causal. For example, decreasing net diversification rates on older islands likely stem from heightened biotic (e.g., increased competition) or abiotic (e.g., rain shadow loss) stressors, rather than island age itself. Incorporating additional biotic and abiotic paleofeatures into analyses would be ideal, yet detailed paleoecological and paleoclimatological data are rarely available for most locations, including the Hawaiian archipelago. Moreover, island biogeography theory is often framed around easily measurable proxies, such as island size, distance, and age, rather than intricate ecological or climatic factors that directly govern biodiversity. For *Kadua*, some features individually exert weak effects on diversification, while the combined effects across features may fundamentally shape long-term evolutionary trajectories. Leveraging larger clades and more complex feature sets will enable future studies to test more nuanced relationships between paleogeographic and biogeographic processes.

We reiterate that Hawaiian *Kadua* is a small clade, so the signal it generates to test biogeographic hypotheses is correspondingly weak. This limited signal is the principal source of the estimation uncertainty we report. Small trees make it difficult to reliably resolve every feature effect across the full range of possible effect strengths, especially weaker ones, which is why we report uncertainty (HPD intervals) and Bayes factors rather than relying on point estimates alone. Our simulations are consistent with this interpretation: for clades comparable in size to Hawaiian *Kadua*, for many it is possible to recover posterior densities and Bayes factors that support signed hypotheses, but entirely excluding the value of zero is harder (“Figures S4-S6”). In addition, false discovery rates for Bayes factors were generally low (“Figure S6”), and posterior estimates of diversification dynamics depart substantially from their priors (“Figures S12-S13”), indicating that the model is not predisposed to affirm biogeographic hypotheses when the appropriate signal is absent.

That the evolutionary history of *Kadua* has been closely intertwined with dynamic paleogeographic changes is not surprising. However, detailed reconstruction of when and where *Kadua* diversified provides a valuable foundation for future studies of insular evolution. Such a framework enables investigations into key shifts in life history, dispersal strategy, reproductive mode, and ecology that make *Kadua* and other island radiations compelling to evolutionary biologists^45^. For example, most *Kadua* lineages exhibit derived secondarily woody growth forms that evolved from herbaceous ancestors, producing diverse trees and shrubs inhabiting moist mid-to high-elevation island forests^23^. *Kadua* also represents one of only 12 documented cases of autochthonous shifts from hermaphroditism to dioecy in the Hawaiian flora, and one of only three such lineages that became species-rich^46^. Morphological and ecological diversity in extant *Kadua* taxa reflects occupation of diverse habitats, from wet forests and bogs to arid, leeward rocky outcrops^47^. By accounting for extrinsic geological influences through our integrative framework, long-standing hypotheses about the drivers and consequences of dispersal limitation, derived woodiness, reproductive transitions, and adaptive radiation can now be robustly tested^45^.

### Unified biogeographic and phylogenetic inference with TimeFIG

To study how life evolved on Earth, biogeographers seek to reconstruct where ancestors of living species originated, dispersed, and went extinct. In practice, this is often done by providing a time-calibrated phylogeny and geographic distribution of extant species to a phylogenetic model, and then inferring the ancestral speciation, extinction, and dispersal events. Although larger phylogenomic datasets have improved the ability to accurately infer species relationships, estimating divergence times remains notoriously difficult for two reasons. First, standard phylogenetic models of molecular evolution cannot separately identify evolutionary rates from geological timescales, so extrinsic information is needed for time-calibration, regardless of genetic sequence data quantity^48,49^. Second, the fossil record remains the primary source of evidence for dating, yet many clades lack suitable fossils. The reliance on fossils to obtain time-calibrated phylogenies, combined with most biogeographic models requiring pre-computed time-calibrated phylogenies as input, means that the historical biogeography of many clades – particularly among plant and insect lineages – remains understudied.

To help overcome these challenges, we developed TimeFIG to jointly infer phylogenetic and biogeographic histories within a paleogeographic context. Our modeling strategy contrasts with alternatives that take a pre-computed time-calibrated phylogeny (or distribution of phylogenies) as input, or do not consider phylogeny whatsoever, to estimate biogeographic rates^15,16,18,50–53^. When computing likelihoods, TimeFIG primarily considers three complementary sources of information: the (co-estimated) phylogenetic relationships and branching times, the presence and absence of each species in each region, and the time-varying paleogeographic features of the regions themselves. For comparison, a standard time-varying GeoSSE model of comparable complexity would only consider the first two data types (a fixed phylogeny and species ranges), and require hundreds of parameters to model the Hawaiian system. The TimeFIG model in this study, in contrast, uses just 20 parameters to infer time-varying and paleogeographically informed rates, with reversible-jump Markov chain Monte Carlo (RJMCMC) effectively reducing this count further. As a generative model, TimeFIG can simulate data under alternative evolutionary scenarios to measure its expected performance in real world scenarios (“Figures S1-S6”). Our simulations show TimeFIG is trustworthy when detecting effects of paleoregional features on biogeographic event rates, indicating the model is not hopelessly overparametrized for use with small clades resembling Hawaiian *Kadua*.

TimeFIG treats inferences as posterior densities of highly-probable data-generating conditions and scenarios, mitigating problems related to an over-reliance on point estimates for weakly or non-identifiable parameters^49,54^. Relatedly, the structure of TimeFIG helps constrain rate estimates that can be difficult to identify in other settings. Non-state-dependent diversification models, for example, allowing speciation and extinction rates to vary freely through time can fit many distinct rate trajectories to an extant-only phylogeny equally well^55,56^. In contrast, TimeFIG rate trajectories do not vary freely, and instead are structured by species tip states (i.e., ranges), temporally varying paleoregional features, and time-constant feature-dependent effect parameters. This added structure does not eliminate estimation uncertainty, but instead limits the histories the model entertains to those compatible with the observed ranges and the external paleogeographic record^57^, i.e. ancestral species cannot disperse into or speciate upon submerged islands. New theory further suggests that certain diversification models with cladogenetic state changes (as TimeFIG has) do not suffer from rate-time non-identifiability^58^. These qualities of TimeFIG make it ideal for for modeling phylogenetic biogeography within paleogeographic systems, where eliminating estimation uncertainty is virtually impossible.

Regardless of how it is done, jointly modeling phylogeny, biogeography, and paleogeography helps unify ecological and evolutionary perspectives on biotic assembly^59^. Classic ecological models such as the equilibrium theory of island biogeography^7^ focus on the balance of biogeographic processes, like immigration and extinction, over shallower timescales to explain species accumulation, rather than the evolutionary processes, like speciation, that operate over deep time. When enhanced for deeper timescales, island biogeography models generally treat species as interchangeable, and do not make full use of information regarding the particular phylogenetic relationships, origination times, and biogeographic histories of ancestral species^8,18^. Conversely, phylogenetic models of biogeography^15,16,51^ typically require a time-calibrated tree is provided before the analyses, and center on evolutionary processes without explicit representations for how paleogeographic forces shape the tree-generating process or modulate ecological species richness. Continued work in this area will hopefully lead to a convergence between the fields of island biogeography and phylogenetic biogeography^8,18,51,60,61^.

## CONCLUSIONS

In this study, we introduced a Bayesian inference framework that leverages paleogeographical dynamics to co-estimate time-calibrated phylogenetic and biogeographic histories. Our analysis of Hawaiian *Kadua* provides empirical support for the influence of island distances and ages on dispersal patterns, and is consistent with the general dynamic theory of oceanic island biogeography in that net diversification rates appear to track island ontogeny, rising during periods of island uplift and habitat expansion, then declining as islands erode. Inferring island diversification without simultaneously considering phylogeny, biogeography, and paleogeography risks missing key interactions between evolutionary history and spatiotemporal context. The re-examination of island radiations through an integrative lens will deepen and refine our understanding of the general principles of evolution and biogeography.

## RESOURCE AVAILABILITY

### Lead contact

Requests for further information and resources should be directed to and will be fulfilled by the lead contact, Isaac H. Lichter-Marck.

### Materials availability

This study did not generate new materials

### Data and code availability

- The raw sequencing files are available at the NCBI database under accession XXXXXXX (to be completed upon acceptance of the article).
- The assembled loci used for this study are hosted at https://github.com/Hawaiian-plant-biogeography/kadua_timefig.
- The scripts used for this study are hosted at https://github.com/Hawaiian-plant-biogeography/kadua_timefig.

## ACKNOWLEDGMENTS

We are grateful to the staff of herbaria that generously provided tissue samples for this study (BISH, PTBG, US), and to Dave Lorence helping us develop our taxon sampling strategy. We thank Sonal Singhal, Maria José Sanín, and participants of the Phylogenetic Biogeography Workshops for providing feedback on earlier versions of the manuscript. We thank Shiva Nia, Anthony Baniaga, Ioana Anghel, Sara Sofía Pedraza, Athena Lam, Sarah Jacobs, Ricardo Kriebel, Ixchel González-Ramírez, Sean McHugh, Sophia Winitsky, Louise Winther, Tim Flynn, Ben Nyberg, Seana Walsch, Dustin Wolkis, Susan Fawcett for helpful discussions. This research was supported by the US National Science Foundation (NS) DEB 2040347 to MJL, 2040081 to FZ, 2040097 to WW, and DBI 2209393 to ILM.

## AUTHOR CONTRIBUTIONS

Conceptualization: I.H.L.-M., S.K.S., F.K.M., F.Z., M.J.L. Investigation: I.H.L.-M., S.K.S., F.K.M., B.G.B., K.R.W., N.R., W.L.W., F.Z., MJL. Writing original draft: I.H.L.-M., S.K.S., F.K.M., F.Z., M.J.L. Writing and editing final draft: all authors. Funding acquisition: W.L.W., F.Z., M.J.L.

## SUPPLEMENTAL INFORMATION INDEX

Document S1. Supplemental methods, Figures S1-S13, and Tables S1-S12

Data S1. Spreadsheet of analyzed *Kadua* samples and GenBank accessions (hosted on GitHub)

## STAR METHODS

### Key resources table

See attached document.

### Experimental model and study participant details

Our sampling includes 23 representatives of 30 currently recognized species of *Kadua* Cham. & Schltdl. and is the most densely sampled phylogenetic study to date of the Hawaiian *Kadua* radiation. We follow the taxonomic checklist on the Flora of Hawaii project website hosted by the Smithsonian Museum of Natural History^62^, although we note that one taxon, *Kadua fosbergii* (W. L. Wagner & D.R. Herbst) W. L. Wagner & Lorence has recently been recognized as *Kadua munroi* (Fosberg) Govaerts in preparation for the Plants of the World Online project^63^. We sample all taxa of *Kadua* in Hawaii with the exception of the rare *Kadua coriaceae* (J. E. Smith) W. L. Wagner & Lorence or the extinct *K. foliosa* Hillebr. and *K. degeneri* subsp. *coprosmifolia* (Fosberg) W. L. Wagner & Lorence, for which DNA extracts were impractical to obtain. In addition, we sampled three taxa of *Kadua* sect. *Austrokadua* (https://www.ipni.org/a/18913-1 Fosberg) W. L. Wagner & Lorence from French Polynesia, *K. lichtlei* Lorence & W. L. Wagner*, K. nukuhivensis* (J. Florence & Lorence) W.L.Wagner & Lorence, and *K. rapensis* F. Br. We further integrated our dataset with publicly available data from the GenBank Sequence Read Archive (SRA) for five outgroups *Exallage auricularia* (L.) Bremek.*, Hedyotis ceylanica* N. Wikstr. & Neupane*, Hedyotis* L.*sp., Scleromitrion polycladum* (F. Meull.) K.L. Gibbons, and *Synaptantha tillaeacea* Hook.F. For *Kadua* found in the Hawaiian islands, we included multiple conspecific samples for each recognized taxon to test the monophyly of current species-level classifications for *Kadua* and uncover potential cryptic diversity. Specifically, we included one sample for each island for each taxon for widespread lineages and two conspecific samples for single island endemic taxa from geographically distant locations, such as distinct valleys or volcanoes. A comprehensive list of samples included and associated data, including GenBank accession numbers is presented in Data S1.

## Method details

### DNA Extraction, Library Preparation, and Sequence Assembly

We extracted DNA from pulverized herbarium and silica dried leaf tissues using the modified CTAB-extraction approach^64^ developed for degraded herbarium tissues described in detail in Johnson et al. (2023)^65^. After destructive sampling from herbarium specimens or silica-dried field collections, leaf fragments were immersed with 96 percent ethanol. Ethanol was evaporated off of leaf tissues during 24 hours of incubation in a fume hood while sample tubes were uncapped and covered with a kim wipe (Kimberly-Clark, Irving, Texas, USA). Following ethanol pretreatment, samples were homogenized in a MillMix 20 Bead Beater (Domel, Zeleznicki, Slovenia) and each homogenized plant powder was suspended in a lysis buffer that consisted of 500µL of SDS (0.5 percent SDS, 0.08 M NaCl, 0.16 M sucrose, 0.064 M EDTA, 0.12 M Tris base, pH 9.1), 10µL of 50 mg/mL proteinase K and 10µL of *β*-mercaptoethanol and then incubated in a Benchmark BSH1002 dry bath incubator (Marshall Scientific, Hampton, New Hampshire, USA) at 54°C overnight. Lysed tissues were purified and DNA was isolated using a chloroform based extraction, in which samples were combined with 500 µL of 24:1 chloroform:isoamyl alcohol and mixed for 3 min. Following centrifugation for 15 minutes, the upper aqueous phase was transferred to a new sample tube, mixed with a 0.08X volume of 7.5M ammonium acetate and a 0.54X volume of isopropyl alcohol, and incubated overnight. After centrifuging these mixtures to pelletize precipitated DNA, we washed extracts twice with 1mL of ethanol wash buffer (80 percent ethanol, 0.02M NaCl, 02M Tris-HCl, pH 8.0), and resuspended DNA in 100µL of TE buffer (0.01M Tris-HCl, 01M EDTA, pH 8.0). DNA concentrations were determined using a Qubit dsDNA broad range assay kit (Thermo Fisher Scientific, Waltham, Massachusetts, USA), according to the manufacturer’s specifications.

For each sample, 400ng of DNA was sent to Daicel Arbor Biosciences (Ann Arbor, Michigan, USA) for quality control, library preparation, target capture, and sequencing. Dual-indexed illumina compatible libraries were prepared by targeting an insert size of 500 bp, with an additional blunt-end repair step for samples deemed of lower quality due to the presence of low molecular weight gDNA. Prior to target enrichment, 6-12 libraries were pooled and each capture pool was dried down to 7 *µ*l by vacuum centrifugation. Captures were performed following the myBaits v5.02 protocol using the Angiosperms353 bait set with an overnight hybridization at 65°C. Postcapture, reactions were amplified using universal Illumina primers and quantified again with a spectrofluorimetric assay. Samples were sequenced on the Illumina NovaSeq 6000 platform on a partial S4 PE150 lane to approximately 1 Gbp per library.

The quality of the demultiplexed raw sequence data was assessed using fastqc^66^. Adapters and low-quality reads were trimmed using Trimmomatic v0.39 with a sliding window of 4 base pairs, cutting when the threshold of average base quality fell below 15 and enforcing a minimum sequence read length of 40 bp^67^. Consensus low-copy nuclear loci were assembled using iterative based reference guided assembly in Hybpiper v.1.3^68^ using the “-bwa” option to match reads to loci. For enhanced sequence recovery we used the expanded mega353 target file as a reference^69^. Besides assembling the exonic regions targeted directly by the angiosperms353 baits, we also assembled flanking intronic regions using the intronerate.py script in HybPiper and putative paralogs using the “paralog investigator.py” script. Potentially problematic sequences were removed from subsequent analyses by concatenating the regions listed in the “genes with long “paralogs.txt” and “genes derived from putative chimeric stitched contigs.txt” output files and using a custom script to remove these regions from each sample in which they were flagged. A modified version of the hybpiper “stats.py” script was then used to generate a gene recovery report and a heatmap that allowed for the filtering of low-performing samples, which we defined as those with in the bottom 10 percent of capture success of orthologous loci across our samples. In addition, we removed samples with less than 1 GB of initial raw sequence data. Twelve samples were excluded from further analyses based on these criteria, resulting in a final dataset of 64 samples. Following sensitivity analyses to determine appropriate data subsets that would maximizing robust statistical support while minimizing potential sources of noise from missing data, we based our final phylogenetic analyses on supercontigs, which included both exons and their more variable flanking introns. Prior to phylogenetic analyses, all loci were aligned using Mafft^70^ under default settings and trimmed using the utility trimal^71^ to remove all sites that exceeded a gap threshold of 80 percent. Our final aligned data matrix included 343,433 bp and 75,658 parsimony informative sites from 344 loci. Average gene and gene tree properties across loci of our dataset produced in downstream analyses are summarized in Table S1.

### Phylogenetics

We initially inferred phylogenetic trees based on a concatenated data matrix using maximum likelihood in IQ-TREE 2^72^. Model selection for each locus was carried out in IQ-TREE 2 using edge-linked proportional models compared with Bayesian Information Criterion scores. A maximum likelihood phylogenetic tree was then inferred based on the partitioned data matrix and support estimated during 1000 ultrafast bootstrap replicates. For each internal branch, gene concordance factors (gCF) were calculated as the fraction of gene trees concordant with that branch and site concordance factors (sCF) as the fraction of informative alignment sites in support of that branch.

In addition to this concatenated analysis, we also independently generated gene trees for each locus using the same procedure in IQ-TREE 2 and used these data to infer a species tree while accounting for incomplete lineage sorting in ASTRAL-III^73^. Prior to their use in ASTRAL-III, low support branches in gene trees where bootstrap support values were less than 20 were collapsed. To visualize phylogenetic trees we used FigTree^74^ and the R package phytools^75^.

Initial phylogenetic trees were reviewed based on expert familiarity with *Kadua* species morphology, ecology, and systematics. Herbarium specimens of samples identified as having anomalous phylogenetic relationships were either examined for misidentification, identified as putatively undescribed taxa, or flagged as potentially contaminated and removed from further analyses. The determination of one sample as *Kadua centranthoides* Hook. & Arn. (SKW 414) was found to be an error and the identity updated to *K. foggiana* (Fosberg) W. L. Wagner & Lorence. The widespread *K. cordata* Cham. & Schtdl. species complex was found to be non-monophyletic, although this complex contains two distinctive infraspecific taxon *K. cordata* subsp. *remyi* (Hillebr.) W. L. Wagner & Lorence and *K. cordata* subsp. waimeae (Wawra) W. L. Wagner & Lorence that were supported as natural groups. After detailed examinination of one sample initially determined as *Kadua cordata* subsp. *cordata* from Molokai (Wood 14018) found to be non-monophyletic with other *K. cordata*, it was determined to be an undescribed taxon in need of further study. One sample of *K. cookiana* Cham. & Schltdl. (Perlman 24732) was resolved with high support as sister to a conspecific sample (Lorence 8817) of the same species but on an exceptionally long branch. To prevent branch length bias introduced by this outlier sample into dating estimates, we excluded it from further analyses.

Prior to introducing our phylogenetic tree for *Kadua* into the TimeFIG modeling framework, we reduced our sample down to one representative per minimum rank taxon, except in cases where taxa were found to be non-monophyletic. Thus, for *K. cordata* we retained four separate operational taxonomic units to accommodate previously unrecognized taxonomic diversity in this species complex. Representative samples were chosen in consultation with experts in *Kadua* natural history to ensure they conformed morphologically and geographically with canonical taxonomic concepts and did not represent outlier samples that might bias analyses. Alignments were reduced using the subset list of samples and a phylogeny for these samples was inferred using ASTRAL, following the procedure described above. We converted the ASTRAL phylogeny to a chronogram using penalized-likelihood with the chronos function in the R package phytools^75^. This chronogram was later provided to RevBayes to initiate RJMCMC analyses using a secondary node calibration for the divergence of the *Scleromitrion*+*Exallage*+*Hedyotis sp.* out-group clade from the *Kadua* ingroup at 20.6 [15.9, 25.3] Ma^23^.

We also selected a subset of 10 loci out of the full set of 344 low copy nuclear loci used in phylogenetic analyses to inform a molecular substitution model for TimeFIG analysis. Our dataset was subset by assessing each of 344 supercontigs using statistical tools included in the *genesortR* script^76^, which summarizes a variety of informative gene and gene tree properties for each locus (“Table S1”). We used the principal component analysis output provided by *genesortR* to identify and avoid outlier loci and sort all others based on their utility as representatives of the overall properties of the full dataset.

### Region definitions

The Hawaiian archipelago contains over one hundred islands, islets, and atolls, eight of which comprise the modern high islands: Hawai‘i, Maui, Moloka‘i, Lana‘i, Kaho‘olawe, O‘ahu, Kaua‘i, Ni‘ihau. We defined six island regions that include the four modern high island complexes where all Hawaiian *Kadua* live, plus two older island complexes that are potentially relevant to the biogeographic history of the clade (shown in Figure 2F of main text). The islands of Hawai‘i and O‘ahu were each treated as separate regions. The Maui Nui island complex includes Maui, Moloka‘i, Lana‘i, and Kaho‘olawe, as well as minor coastal islets that have served in some cases as essential reservoirs of botanical diversity, such as Huelo islet off the cliffy north coast of Moloka‘i. The islets of Ni‘ihau and Lehua were likewise included with a broadly defined Kaua‘i island complex. The Necker region encompasses the volcanic remnant islands of Necker, Nihoa and Kalau. We defined the Gardner island complex to include the older, now eroded volcanic islands Maro through the French Frigate Shoals. The seventh outgroup region represents all locations outside of the remaining six Hawaiian regions.

### *Kadua* species ranges

Discrete biogeographical ranges for each taxon were determined based on continually updated geographic distribution data on the Flora of Hawaii project website, hosted by the Smithsonian Museum of Natural History^62^, and from extensive knowledge of field and herbarium collections by co-author Kenneth Wood. Taxa were scored for presence or absence in each of 6 major island complexes, Hawai‘i (H), Maui Nui (M), O‘ahu (O), Kaua‘i (K), Necker (N), and Gardner (G)^1^ or, if it was an outgroup taxon, it was coded as present in only the outgroup region (Z). At an advanced stage of the writing of this manuscript, we discovered two minor discrepancies in the ranges of *Kadua littoralis* and the *Kadua cordata* species complex. For *K. littoralis*, we coded it as present in K, O, and M but absent in H, yet there are reports of its presence in H. For the *K. cordata* species complex, we discovered this is not a monophyletic group, which is recognized with three operational taxonomic units *Kadua cordata* subsp. *cordata*, *Kadua cordata* subsp. *remyi*, and *Kadua cordata* subsp. *waimeae*. The Flora of Hawaii lists the distributional range of *Kadua cordata* subsp. *cordata* as being found on K, O, and M; however, given that we suspect these island populations to constitute cryptic taxa, our dataset assigns this taxon a restrictive range of being found only on M, where it is sympatric with the other two subspecies. Fixing either of these minor discrepancies is not expected to alter our results because both taxa are nested within the clade *Kadua* sect. *Wiegmannia*, where they are closely related to other widespread lineages.

### Paleogeographic epochs

Our model of paleogeography was based on^18,22,27,77^ and includes 7 sequential time intervals (Ma): 32 to 19.7, 19.7 to 10.6, 10.6 to 6.2, 6.15 to 4.15, 4.15 to 2.55, 2.55 to 1.20, 1.20 to 0.0. We set ages equal to the emergence of the oldest island in each island complex. For modern island complexes Hawai‘i, Maui Nui, O‘ahu, and Kaua‘i, we used the ages in^18^. We used 11.5 Ma (mean of 11 and 12 Ma) for the origin time of Necker to represent the Necker complex^77^. We used 20.5 Ma (mean of 20 to 21 Ma), which corresponds to the origin time of Maro, to represent the age of the Gardner complex. All times are listed in “Table S2”.

### Paleogeographic features

To model change among the relevant geographic features over time, and to test the specific hypotheses outlined in the main text, we gathered paleogeographic data for 8 features for each of the 7 regions and 7 time intervals, marked by the appearance times of new island complexes (“Table S3”). All variables for features were either categorical or quantitative, and either described within-region features of singular island complexes or pair-wise relationships between regions.

To test the size-diversification relationship (H1), we modeled island size through time as a quantitative within-region variable based on paleoelevations. Paleoelevation for each region *i* and epoch *k* was computed as a time– and island-averaged mean using the data table of Price et al.^27^. Paleoelevation in^27^ is recorded as *y_ab_* where *a* (row) corresponds to individual islands and *b* (column) corresponds to 0.5 Myr time bins. Entries for *y_ab_* are only positive when islands are above sea level. We represented the paleoelevation for region *i* and epoch *k* as the arithmetic mean of the set of all positive paleoelevations for islands (rows) and 0.5 Myr time bins (columns) corresponding to region *i* and epoch *k*.

To test the distance-dispersal relationship (H2), we measured quantitative inter-island distances (km) using Google Earth. Hawaiian hotspot dynamics have been inconsistent through time, resulting in non-uniform distances between sequential island complexes^11,77^. However, collective rifting of islands along the North American plate means that inter-island distances have changed little through time. We therefore used constant inter-island distances across all time intervals. To infer rate and effects of long distance, trans-oceanic dispersal, we included an additional categorical variable differentiating between dispersal in/out of the archipelago versus dispersal within the archipelago.

To test the progression rule of island biogeography (H3) we categorized island relationships as older to younger or younger to older. We also used midpoint ages for each island complex in each time interval to model the effects of the age of different complexes, and used inter-island age differences (old age minus young age) to capture whether the magnitude and/or sign of island age difference mattered.

To test the predictions of dynamic equilibrium theory (H4), we modeled island growth as a step function using two overlapping categorical within-region variables. The first categorized islands as high islands or not, with high islands defined as having a paleoelevation greater than 1000m^27^. A second categorical variable identifies whether each island complex in each time interval is in a state of net growth or net decay. Together, these categorical variables capture when an island is young (high + growth), middle-aged (high + no-growth), or old (no-high + no-growth).

All quantitative features for all regions were using a signed log-transform before being included into the TimeFIG modeling framework. To remove the influence of the external region (Z), we set each of its feature values to the corresponding mean for the remaining insular regions.

## Quantification and statistical analysis

### TimeFIG model definition

TimeFIG is an extension of the Feature-Informed GeoSSE model^16,41,53^. TimeFIG is a stochastic branching process that models dispersal (d), extinction (e), within-region speciation (w), and between-region speciation (b) events with region-dependent rates that respond to the regional paleogeographic features of the system at time *t*. These events cause species ranges, which are encoded as sets of occupied regions, to evolve during and in-between speciation events. In particular, within-region (*in situ*) speciation occurs at rate *r_w_*(*i, t*), forming a new species in region *i*. Extinction (range contraction) occurs at rate *r_e_*(*i, t*), and causes a species to lose region *i* from its range; any species losing the last region in its range goes globally extinct. Dispersal (range expansion) from region *i* into region *j* occurs at rate *r_d_*(*i, j, t*). Between-region speciation splits an ancestral range *u* ∪ *v* into daughter ranges *u* and *v* at rate *r_b_*(*u, v, t*).

All TimeFIG event rates are defined in a similar manner. Rates for each of the four event types, *p* ∈ {*d, e, w, b*}, are computed as the product of an absolute base rate, *ρ_p_*, and a relative rate modifying factor, *m_p_*, that depends on geographic information stored in the relevant regions at geological time, *t*. Taking within-region speciation rate in region *i* as an example, we have

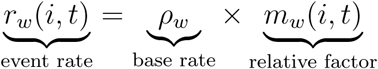

where the within-region speciation rate factor is further defined as

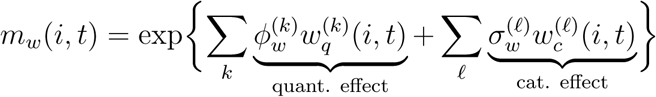

which produces a positive value representing the net effect of all quantitative and categorical (0 or 1) features upon within-region speciation for region *i* at time *t*. Each quantitative or categorical feature layer (*k* or *ℓ*) records regional attributes for each region *i* and time *t*. The feature effect for each layer is the product of an estimated feature effect parameter (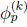 or 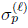) and its corresponding vector of inputted layer-features (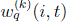 and 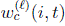).

Note that the base rate (*ρ_w_*) is constant while the relative factors (*m_w_*(*i, t*)) are functions of region (*i*) and time (*t*). The feature effect parameters, 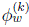 and 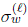, are estimated parameters that do not depend on time or region, whereas the layer-features, 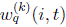 and 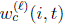, are input values that vary across regions and times. Together, this means that species have a constant relationship with different geographic features, so changes in paleogeographic features induce changes in rates. A total of 20 parameters (4 base rates, 16 feature effect parameters) are used to model the system of time-dependent and region-dependent rates for the empirical analysis.

Each estimated feature effect parameter (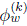 or 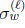) can take any value. A positive feature effect parameter value means the relative rate increases when the feature value is large (e.g., within-region speciation increases with region size). A negative feature effect parameter value means the rate decreases with larger feature values (e.g., dispersal rates decrease as distance increases). A feature effect parameter value of zero means the rate is unaffected by the relevant feature. We use reversible-jump Markov chain Monte Carlo (RJMCMC)^78^ to explore all models with different combinations of feature effect parameters turned “on” (≠ 0) or “off” (= 0). RJMCMC both reduces the effective number of model parameters, and produces posterior estimates that can be used to compute Bayes factors.

We reiterate that all event rates for within-region speciation, *r_w_*(*i, t*), extinction, *r_e_*(*i, t*), dispersal, *r_d_*(*i, j, t*), and between-region speciation, *r_b_*(*u, v, t*) are handled in a similar manner. Relationships between TimeFIG feature effect parameter values and our four biogeographic hypotheses are shown in “Figure 1E” and summarized in “Table S4”. Additional background and details for TimeFIG are provided in the Supplement.

### TimeFIG validation

We used three protocols to validate the behavior of TimeFIG models and the correctness of their implementation in RevBayes and TensorPhylo. See the Supplement for more details.

The first analysis was a Bayesian coverage experiment^79^ under a scenario with two regions, two epochs, one set of feature layers, and a fixed tree (“Figure S2 and Table S6”). We set the target coverage level at 95%, meaning we expect 95% of HPD95 intervals to contain the true data-generating parameter for when the simulating and inference models match. We simulated 100 datasets using PhyloJunction^80^ then estimated posteriors for each dataset using RevBayes and TensorPhylo.

The second analysis simulated island radiation data radiations to measure divergence time estimation accuracy for TimeFIG (“Figure S3”). Each simulation contained the colonization of a single lineage into the Hawaiian islands into a random subaerial island at a random time, separated into scenarios representing either recent colonization between 1 and 6 Ma, or an ancient colonization between 6 and 20 Ma. We used PhyloJunction^80^ to simulate an island “ingroup” radiation under a TimeFIG model that produced realistic biogeographic distributions resembling Hawaiian *Kadua*. Then, we simulated a random crown age and grafted a non-Hawaiian “out-group” lineage to the phylogeny as sister to the ingroup, and simulated molecular sequence data for the full clade. We then estimated posterior densities and compared island clade age estimates to the truth under two inference models.

The third analysis measured how reliably TimeFIG could test biogeographic hypotheses using credible intervals (“Figure S4-S5”) and/or Bayes factors for small clades (“Figure S6”). To do so, we simulated 150 Hawaiian radiations resembling Hawaiian *Kadua*. Each simulation used a different random set of feature effect parameters set to be “on” (value of ≠ 0), and thus involved in diversification or dispersal, or “off” (value of 0), and thus irrelevant for those processes. We then performed Bayesian inference for each simulated dataset, while fixing the tree to the truth, to measure the proportions of intervals that contained the correct value and/or sign of the hypothesis, and Bayes factors supporting each signed hypothesis.

### Empirical Model and RJMCMC settings

We performed joint Bayesian inference of biogeographic rates, phylogenetic divergence times, and molecular evolution under a relaxed clock model using RevBayes^81^ and the TensorPhylo plugin^82^. Using RJMCMC^78,83^, each feature effect parameter (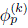 and 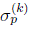) had 0.5 prior probability of no effect (=0) and 0.5 prior probability of following i.i.d. standard normal densities, bounded to [-4, 4]. Other model priors were chosen to be weakly informative and to induce a relatively flat root age density under the prior and uncalibrated analysis. We used 10 high-quality loci and the fixed ASTRAL topology to reduce the runtime of the analysis. The main text presents results for three of eight model settings we explored (M1: TimeFIG calibration, M5: node calibration, M7: no calibration; “Table S5”). Multiple independent MCMCs for each model setting yielded very similar posterior distributions for key model quantities. We found no indication that different chains were isolated to different posterior modes, so we combined MCMC chains to ensure 200+ ESS for all key parameters and quantities, and to improve the accuracy of HPD interval estimates. See the Supplement for more details.

### Software

We used RevBayes^81^ for Bayesian model design and inference along with TensorPhylo^82^ to accelerate the computation of SSE model likelihoods. We added a new family of classes to RevBayes to help users input and manage time-varying regional feature data for FIG model composition. We also modified TensorPhylo to allow the tree variable to be estimated (not fixed) during inference, with efficient sharing of unchanged partial likelihoods during the exploration of tree space. We also improved the stability of TensorPhylo when its numerical integrators are tasked with solving difficult systems of differential equations, like those that arose for this study (e.g., extreme rate variation relative to states, time intervals, and branch lengths). See the Supplement for more details.

### Additional resources

Tutorials describing the theory and application of TimeFIG to *Kadua* are available at https://revbayes.github.io/tutorials/fig_intro/

