## Supplementary Information for "Synchronizing speciation, extinction, and dispersal to island paleodynamics through Bayesian phylogenetics in a Hawaiian plant radiation"

### TimeFIG model design

#### Overview

TimeFIG is a time-heterogeneous feature-informed GeoSSE (FIG) model that describes how species diversify among paleogeographic regions over time. As a member of the GeoSSE model family [4], TimeFIG is a state-dependent stochastic branching process with four major event types: dispersal, extinction, within-region speciation, and between-region speciation. The exact rates for these events depend on the model parameters, the set of regions currently occupied by a species (its range), the time-calibrated paleogeographic features of region(s) relevant biogeographic event, and the model parameter values. We use $w$, $e$, $d$, and $b$ as labels to denote within-region speciation, extinction, dispersal, and between-region speciation, respectively.

Within-region speciation events occur in region $i$ at time $t$ at rate $r_w(i,t)$, and model a new daughter species originating within one region of the parental species range, while the other species retains the parental range. For example, within-region speciation acting on parental species with range $AB$ could yield a new daughter species with range $A$ or $B$ while the other daughter fully inherits (retains) $AB$.

Extinction events occur in region $i$ at time $t$ at rate $r_e(i,t)$, and cause the species to lose one region from its range. For example, extinction will cause the species with range $AB$ to contract to a range of just $A$ or $B$. Losing the last region from a range causes lineage-level extinction, meaning a species in only region $A$ that experiences extinction must lose $A$ and is removed from the pool of surviving species.

Dispersal events from region $i$ into $j$ at time $t$ occur at rate $r_d(i,j,t)$, causing a species to expand its range by a new region. For example, a species with range $AB$ may gain region $C$ either through dispersal of $A$ to $C$ or $B$ to $C$.

Between-region speciation events split a widespread species (i.e., occupying two or more regions) with range $u \cup v$ into two new species with ranges $u$ and $v$, such that $u \cap v = \emptyset$, at rate $r_b(u,v,t)$. For example, the widespread species $ABC$ may yield pairs of daughter species with ranges $AB$ and $C$, $AC$ and $B$, or $BC$ and $A$.

There are three ways to group the four events. The first way is by anagenetic versus cladogenetic events, where extinction and dispersal occur along phylogenetic branches (anagenetic events), whereas within-region and between-region speciation create new branches (cladogenetic events). The second way is by within- versus between-region events, extinction and within-region speciation occur within individual regions, whereas dispersal and between-region speciation occur between regions. Lastly, in terms of species richness, dispersal and within-region speciation always increase regional species richness, whereas between-region speciation and extinction either conserve or reduce regional species richness.

#### Event rates

TimeFIG event rates for process $p \in \{d,e,w,b\}$ are computed as the product of an absolute base rate, $\rho_p$, and a relative rate modifying factor, $m_p$, that depends on geographic information stored in region $i$ (and possibly a second region, $j$) and geological time, $t$. All TimeFIG rates are defined in a similar manner.

Taking within-region speciation rate in region $i$ as an example, we have

$$\underbrace{r_w(i,t)}_{\text{event rate}} = \underbrace{\rho_w}_{\text{base rate}} \times \underbrace{m_w(i,t)}_{\text{relative factor}} \tag{1}$$

where the within-region speciation rate factor is further defined as

$$m_w(i,t) = \exp\left\{ \sum_k \underbrace{\phi_w^{(k)} w_q^{(k)}(i,t)}_{\text{quant. effect}} + \sum_\ell \underbrace{\sigma_w^{(\ell)} w_c^{(\ell)}(i,t)}_{\text{cat. effect}} \right\} \tag{2}$$

which produces a positive value representing the net effect of all quantitative and categorical features upon within-region speciation for region $i$ at time $t$. Each quantitative or categorical feature layer ($k$ or $\ell$) records regional attributes for each region $i$ and time $t$. The feature effect for each layer is the product of an estimated feature effect parameter ($\phi_p^{(k)}$ or $\sigma_p^{(\ell)}$) and its corresponding vector of inputted features ($w_q^{(k)}(i,t)$ and $w_c^{(\ell)}(i,t)$).

Our main analyses used natural log-transformed measurements for quantitative features. The relative rate effect contributed by natural log-transformed features in Equation 2 simplifies as

$$\begin{aligned} m_w(i,t) &= \exp\left\{ \sum_k \phi_w^{(k)} \ln w_q^{(k)}(i,t) \right\} \\ &= \prod_k \exp\left\{ \phi_w^{(k)} \ln w_q^{(k)}(i,t) \right\} \\ &= \prod_k \left[ w_q^{(k)}(i,t) \right]^{\phi_w^{(k)}} . \end{aligned}$$

Our experience with TimeFIG is that log-transformed quantitative features allow for weaker and subtler relationships between regional features and biogeographic rates, that are both easier to estimate and more realistic in effect sizes.

Note that in Equation 1, the base rate ($\rho_w$) is constant while the relative factors ($m_w(i,t)$) are functions of time ($t$) and region ($i$). The feature effect parameters, $\phi_w^{(k)}$ and $\sigma_w^{(\ell)}$, are estimated parameters that do not depend on time or region, whereas the layer-features, $w_q^{(k)}(i,t)$ and $w_c^{(\ell)}(i,t)$, are input values that vary across regions and times. Each estimated feature effect parameter ($\phi_w^{(k)}$ or $\sigma_w^{(\ell)}$) can take any value. A positive feature effect parameter value means the relative rate is large when the feature value is large (e.g. within-region speciation increases with region size). A negative value means the rate decreases with larger-valued features (e.g. dispersal rates decrease as distance increases). A zero value means the rate is the same for all regions regardless of the relevant feature. In practice, we use reversible-jump Markov chain Monte Carlo (RJMCMC) to explore all combinations of models with different feature effect parameters turned "On" ($\neq 0$) or "Off" ($= 0$).

All other biogeographic event rates are computed in an analogous manner. Extinction rates in region $i$ at time $t$ are defined as

$$\begin{aligned} r_e(i,t) &= \rho_e \times m_e(i,t) \\ m_e(i,t) &= \exp\left\{ \sum_k \phi_e^{(k)} w_q^{(k)}(i,t) + \sum_\ell \sigma_e^{(\ell)} w_c^{(\ell)}(i,t) \right\} \end{aligned}$$

where $w_q^{(k)}$ and $w_c^{(\ell)}$ describe the same paleogeographic feature layers used to model within-region speciation.

Dispersal rates from region $i$ into $j$ at time $t$ are defined as

$$\begin{aligned} r_d(i,j,t) &= \rho_d \times m_d(i,j,t) \\ m_d(i,j,t) &= \exp\left\{ \sum_k \phi_d^{(k)} b_q^{(k)}(i,j,t) + \sum_\ell \sigma_d^{(\ell)} b_c^{(\ell)}(i,j,t) \right\} \end{aligned}$$

where $b_q^{(k)}$ and $b_c^{(\ell)}$ describe paleogeographic feature layers for all pairs of regions.

Between-region speciation rates that produces daughter species with ranges $u$ and $v$ are defined as

$$\begin{aligned} r_b(u,v,t) &= \rho_b \times f_b(m_b(u,v,t)) \\ m_b(u,v,t) &= \exp\left\{ \sum_k \phi_b^{(k)} b_q^{(k)}(i,j,t) + \sum_\ell \sigma_b^{(\ell)} b_c^{(\ell)}(i,j,t) \right\} \\ f_b(u,v,t,m_b) &= \frac{1}{Z} \times \left[ \sum_{(e_1,e_2)\in E(u,v)} \bar{m}_b(e_1,e_2,t)^{-1} \right]^{-1} \end{aligned}$$

where $f_b$ is the range split score function [10], $E(u,v)$ is the cutset of edge to split ancestral range $u \cap v$ into daughter ranges $u$ and $v$, $\bar{m}_b(e_1,e_2,t)$ is the arithmetic mean $(m_b(e_1,e_2,t) + m_b(e_2,e_1,t))/2$, and $Z$ is the geometric mean of all range split scores in $m_b$. In practice, $f_b$ means that ranges split fastest against those region-pairs $(e_1, e_2)$ that have large values $m_b(e_1,e_2,t)$ at time $t$. Note that $m_b$ uses the same between-region features as $m_d$.

#### Biogeographic hypotheses

One goal of this study is to use TimeFIG parameter estimates to test which of four key biogeographic hypotheses apply to Hawaiian *Kadua*: the size-diversification hypothesis [12], the distance-dispersal hypothesis [12, 1], the progression rule of island biogeography [3], and the general dynamic theory of oceanic island biogeography [20]. We test these hypotheses by considering the signs and magnitudes of different feature effect parameter values governing the relevant process-feature relationships, as summarized in Table S4.

We measure statistical support that a process and feature follow the hypothesized relationship ($H_1$) rather than not ($H_0$) using Bayes factors [18]. The Bayes factor supporting hypothesis $H_1$ over $H_0$ is computed as

$$K = \frac{P(X \mid H_1)}{P(X \mid H_0)} = \frac{P(H_1 \mid X)}{P(H_0 \mid X)} \times \frac{P(H_0)}{P(H_1)} = \frac{P(H_1 \mid X)}{P(H_0 \mid X)} \times \frac{0.75}{0.25}$$

where $P(H_1 \mid X)$ and $P(H_0 \mid X)$ are posterior probabilities from RJMCMC, $P(H_0)$ and $P(H_1)$ are prior probabilities, $H_1$ represents a feature effect value is non-zero and has a sign matching the biogeographic hypothesis, and $H_0$ represents a feature effect value that is zero or has a sign that is opposite of the biogeographic hypothesis. Our RJMCMC assigns equal prior probabilities of 0.5 for any feature effect being on or off. Feature effect parameters that are on follow a prior density of N(0,1), which also assigns equal probabilities of 0.5 to the effect being positive or negative. As we are considering signed hypotheses, $H_1^+$ represents the hypothesis for a positive feature effect parameter value ($> 0$) and $H_0^+$ corresponds to any other relationship ($\leq 0$). Similarly, $H_1^-$ and $H_0^-$ correspond to support and non-support for a negative hypothesized relationship. This means the prior probability supporting $H_1^+$ (or corresp. $H_1^-$) is $P(H_1^+) = 0.5 \times 0.5 = 0.25$, and the probability opposing it is $P(H_0^+) = 1 - 0.25 = 0.75$. We follow Kass & Rafferty [9] to interpret the strength of evidence $K$ offers in support of hypothesis $H_1$ over $H_0$: $< 1.0$ = Opposing, $\approx 1.0$ = None, $> 1.0$ = Marginal, $>3.2$ = Substantial; $> 10$ = Strong, $>32$ = Very strong.

#### Bayesian model design

We used RevBayes to jointly estimate all biogeographic, molecular, and phylogenetic model parameters. This section describes the 'full' model we used for the main results of the study (called model M1). Later sections describe how the M1 design was modified to suit the purposes of other analyses and experiments.

Species ranges and phylogenetic divergence times were modeled using a TimeFIG process. We used separate sets of priors for parameters governing dispersal, extinction, within-region speciation, and between-region speciation. Base biogeographic event rates for $p \in \{d, e, w, b\}$ were defined as $\rho_p \sim \text{Exp}(10)$. Feature effect parameters ($\phi_p^{(k)}$ and $\sigma_p^{(\ell)}$) were modeled with reversible jump distributions, placing 0.5 prior probability on the point mass of 0 (no effect) and 0.5 prior probability on the parameter following the prior Norm(0,1). We truncated the prior to values of $[-4, 4]$ to prevent the need to evaluate unlikely but computationally expensive model likelihoods under extreme parameter values proposed by MCMC. Model variants that conditioned on paleogeographic island ages had root state frequencies of $\pi_i^{bg} = 0$ for all states except $\pi_Z^{bg} = 1$ to force the clade to originate in the external region, $Z$. Model variants that ignored paleogeography assumed a uniform distribution over root states.

Nucleotide sequences for locus $i$ were modeled using an HKY85 substitution process [6] with base frequencies $\pi_i^{mol} \sim \text{Dirichlet}(1)$ and transition-transversion rate-ratio of $\kappa_i \sim \text{Gamma}(2, 2)$ [mean 1] and among site rate variation using a $+\Gamma4$ model with shape and scale parameter $\alpha_i \sim \textit{Exp}(0.1)$. Molecular clock rates for locus $i$ and branch $j$ were defined as $\mu_{ij} = \mu^{base} \times r_i^{locus} \times r_j^{branch}$, with base clock rate $\mu^{base} \sim \text{Exp}(10)$, locus rate $r_i^{locus} \sim \text{Exp}(1)$, and branch rate $r_j^{branch} \sim \text{Lognorm}(ln(\mu^{base}) - 0.5, 1)$ [mean 1]. To remove an unnecessary degree of freedom, we always assume the distribution for $r_1^{locus}$ a point mass at the value 1 (i.e. no effect).

#### Posterior probability of full model

We express the probability of observing biogeographic range data $X^{bg}$ and time-calibrated phylogeny $\Psi^{bg}$ as

$$Pr(X^{bg}, \Psi^{bg} \mid \theta^{bg}, Y^{pg})$$

where $\theta^{bg} = (\pi^{bg}, \rho_d, \rho_e, \rho_w, \rho_b, \phi_d, \phi_e, \phi_w, \phi_b, \sigma_d, \sigma_e, \sigma_w, \sigma_b)$ are the biogeographic model parameters, and $Y^{pg}$ is a fixed variable containing information about all paleogeographic features within and between all regions with respect to geological time.

Next, we define the probability of observing molecular data $X^{mol}$ as the product of independent per-locus probabilities,

$$Pr(X^{mol} \mid \theta^{mol}, \Psi^{bg}) = \prod_i Pr(X_i^{mol} \mid \theta_i^{mol}, \Psi^{bg}),$$

where $\Psi^{bg}$ is the time-calibrated phylogeny, $\theta_i^{mol} = (\pi_i^{mol}, r_i^{locus}, \kappa_i, \alpha_i)$ are the parameters for locus $i$, and $\theta^{mol} = (\mu^{base}, r_1^{branch}, \ldots, r_J^{branch}, \theta_1^{mol}, \ldots, \theta_L^{mol})$ is the parameter vector containing the base clock rate, $J$ branchwise clock rates, and $L$ locus-specific parameters.

The joint model probability of $X^{bg}$ and $X^{mol}$ is therefore

$$Pr(X^{mol}, X^{bg}, \Psi^{bg} \mid \theta^{mol}, \theta^{bg}, Y^{bg}) = Pr(X^{bg}, \Psi \mid \theta^{bg}, Y^{pg}) \prod_i Pr(X_i^{mol} \mid \theta_i^{mol}, \Psi^{bg})$$

which leads to the Bayesian posterior

$$Pr(\theta^{bg}, \theta^{mol}, \Psi^{bg} \mid X^{bg}, X^{mol}, Y^{bg}) = \frac{Pr(X^{bg}, \Psi^{bg} \mid \theta^{bg}, Y^{pg}) Pr(X^{mol} \mid \theta^{mol}, \Psi^{bg}) Pr(\theta^{bg}) Pr(\theta^{mol})}{Pr(X^{bg}, X^{mol})}.$$

We can compute the prior probabilities for biogeographic parameters as

$$Pr(\theta^{bg}) = Pr(\pi^{bg}) \times \left[ \prod_{p \in \{d,e,w,b\}} Pr(\rho_p) \times \left[ \prod_k Pr(\phi_p^{(k)}) \right] \times \left[ \prod_\ell Pr(\sigma_p^{(\ell)}) \right] \right]$$

and prior probabilities for molecular parameters as

$$Pr(\theta^{mol}) = Pr(\mu^{base}) \times \left[ \prod_{i \in \text{loci}} Pr(\pi_i^{mol}) Pr(\alpha_i) Pr(\kappa_i) Pr(r_i^{locus}) \right] \times \prod_{j \in \text{branches}} Pr(r_j^{branch}).$$

The marginal likelihood, $Pr(X^{bg}, X^{mol})$, is an intractable constant term. However, the same marginal likelihood appears in the numerator and denominator of the Metropolis-Hastings ratio used for Bayesian MCMC, which cancels with itself, so we do not need to compute the quantity.

We use reversible jump MCMC to sample across models of different dimensions in proportion to their posterior probabilities [5]. In particular, we sample within a family of nested models where feature effect parameters are turned on (e.g. $\phi_p^{(k)} \sim \text{Norm}(0,1)$) or off (e.g. $\phi_p^{(k)} = 0$). When we propose a jump from a model where one feature effect parameter goes from a zero (off) to a non-zero (on) value, we sample the new value from that feature effect's prior density, causing its prior probability (numerator) and proposal probability (denominator) to cancel in the Metropolis-Hastings-Green ratio. The same logic applies for the reverse move where we propose a jump from a model with a feature effect with a non-zero value (on) to one with a zero value (off). See the Discussion of [5] for details.

#### Model variants

We analyzed eight model variants (M1–M8) to assess how modifying five analysis settings influenced parameter estimates (Table S5). When the "Use data?" setting was false, the model ran the analysis under the prior, meaning molecular and biogeographic data did not inform the likelihood function. When "Use features?" was false, it assumed all feature effect parameters ($\phi$ and $\sigma$) were fixed to zero, meaning all biogeographic rates would equal the base rate ($\rho$) for the corresponding process. When "Use paleogeo.?" was false, it assumed that all islands always existed and had regional features identical to those at present. When "Use 2° calibration?" was true, it applied the Neupane et al. [16] node age calibration to the MRCA of *Exallage* and Hawaiian *Kadua*. When "Use biogeo. calibration?" was true, it applied a maximum clade age constraint on the MRCA of Hawaiian *Kadua*.

##### MCMC analyses

Using RevBayes [8] with the TensorPhylo plugin [13], we performed multiple, independent MCMC analyses for each model variant, with more runs for more computationally intensive variants: 7x runs for M1; 5x runs for M2; 4x runs for M3; 4x runs for M4; 3x runs for M5; 2x runs for M6; 2x runs for M7; and 2x runs for M8. Multiple runs allowed us to verify MCMC convergence and combine independent, convergent runs to increase the accuracy of posterior parameter estimates. For each chain, we removed the first 10% of MCMC samples as burn-in, and then assessed convergence of each chain using Tracer. No MCMC traces contained erratic jumps in posterior probability, nor contained posterior samples that substantially differed from other MCMC traces (e.g. 95% credible intervals from different chains had no overlap). All MCMC traces for each model variant were then combined (without burn-in) into a single trace file, for which the effective sample sizes (ESS) for important parameters was always above 100. Relevant results are discussed in the main text.

#### TimeFIG model assessment

##### Rate-time identifiability

Standard substitution molecular substitution models cannot separately identify rate from time when computing transition probabilities. Any combination of evolutionary rates, $r$ or $r'$, and durations of time, $t$ or $t'$, with rate-time products satisfying $rt = r't'$, that evolve under the same rate matrix $Q$, yield the same transition probability matrix, $P(r,t) = \exp(Qrt) = \exp(Qr't')$, and assign identical probabilities to evolutionary outcomes. We refer to this as rate-time non-identifiability.

Biogeographic dating with TimeFIG makes use of the fact that the model's event rates vary across states (ranges) in response to paleogeographic events that are calibrated to geological time. As a result, the TimeFIG model transition probabilities will vary as a function of time in response to the modeled paleogeographical scenario. To demonstrate this, we take an arbitrary example of the *Kadua* timetree to measure how its likelihood changes as a function of tree height.

To test for biogeographic rate-time identifiability in TimeFIG, we rescale the timescale of the phylogeny, the paleogeographic events, and the units of the event rates, then compare model likelihoods across different model variants (described later). We ignore molecular substitution model likelihoods, since they are already known to have rate-time non-identifiability. To demonstrate rate-time non-identifiability for continuous-time Markov models (e.g., molecular substitution models), one can multiply all branch lengths by a factor of $x$ and the evolutionary rate by a factor of $1/x$, yet produce the same model probability. Changing the timescale, however, changes the scale (but not the shape) of the birth-death model probability density [7]. To control for this sensitivity, rather than rescaling all phylogenetic ages and rates, we instead hold the phylogenetic tree height and rates constant and rescale the relative timing of paleogeographic ages as $\tilde{a}_k^{island} = a_k^{island} \times a^{ingroup}/\tilde{a}^{ingroup}$, where $a_k^{island}$ is the true age of island $k$, $a^{ingroup}$ is the true age of the ingroup clade age, $\tilde{a}^{ingroup}$ is effective rescaled age we want to assign to the ingroup, and $\tilde{a}_k^{island}$ is the effective rescaled age of the islands relative to the tree. Using this transformation, we compute likelihoods under each model variant where the effective ingroup clade age ranged from 0.1 to 29.6 Ma at 0.5 Myr intervals. To better compare model fit between model variants and across timescales, we normalized the raw likelihoods for each model variant by the sums of their respective likelihoods, and used these for display.

We then analyzed the tree using three biogeographic models. The first model assumes an equal-rates GeoSSE model that allows species to occupy islands at any time, even before island formation. This variant is most similar to M7. It does not have time-heterogeneous rates, so we expect it contains no time-calibration information. The second model assumes that we have an equal-rates GeoSSE model that only allows species to occupy islands after formation. This variant is a TimeFIG model with all feature effect parameters set to zero (no effect), and is similar to M3. It has time-heterogeneous rates that only restrict dispersal opportunities, which we expect inform maximum (not minimum) clade ages. The third model assumes the full TimeFIG model, for which island existence and other paleogeographic features jointly influence island occupancy and biogeographic rates. This variant is similar to M1. It has time-heterogeneous feature-dependent rates, which restricts dispersal and creates periods of higher and lower net diversification rates, such as those predicted by Dynamic Island Biogeography Theory [20]. Because of this, we expect the full TimeFIG model will generate information for both maximum and minimum clade ages.

For the three models, we used an arbitrary *Kadua* timetree and model parameters to measure how biogeographic likelihoods change as a function of absolute timescale. We set the biogeographic base rate parameters to $\rho_w = 0.1$, $\rho_e = 0.1$, $\rho_d = 0.05$, and $\rho_b = 0.2$. For the full TimeFIG analysis, we set the feature effect model parameters to emulate island biogeographic dynamics, using the following values: $\phi_d^{dist} = -1, \phi_d^{\Delta age} = 0, \phi_e^{size} = -0.25, \phi_e^{age} = 0.25, \phi_w^{size} = 0.25, \phi_w^{age} = -0.25, \phi_b^{dist} = 1, \phi_b^{\Delta age} = 0, \sigma_d^{dist} = -2, \sigma_d^{\Delta age} = -0.25, \sigma_e^{high} = -0.25, \sigma_e^{grow} = 0.25, \sigma_w^{high} = 0.25, \sigma_w^{grow} = -0.25, \sigma_b^{dist} = 2, \sigma_b^{\Delta age} = 0$.

The relative likelihood curves are shown in Figure S1. Time-constant GeoSSE model probabilities (M7) show no difference in relative likelihoods for different combinations of $rt = r't'$, as expected. Note that a biogeographic node age calibration under (e.g. M5, not shown) would appear similar to M7 (flat) but assign -inf log-likelihoods to all clade ages older than Kauai ($\sim 6.2$ Ma). The model that accounts for paleogeographic island ages but not features (M3) produces log-likelihoods that respond to the time scale, assigning higher scores to ingroup clade ages of between approximately 0 and 15 Ma. Lastly, the TimeFIG model (M1) shows similar dynamics as the previous example, but a narrower window of higher scores between approximately 0 and 6 Ma. M1 and M3 both use paleogeographic island ages, but only M7 uses paleogeographic regional features. The difference in supported ingroup clade ages reflects the fact that M3 allows species to survive indefinitely on older, low islands, whereas M1 assigns higher extinction and lower speciation to lineages on those islands through the regional features.

#### Coverage experiment

We validated a simple TimeFIG model by carrying out a Bayesian coverage validation experiment [14]. Our coverage validation has three steps: simulation, inference, and comparison. The simulation step draws data-generating parameters from the prior distribution of the model (the "true" parameter values) and then simulates one phylogenetic dataset for each random set of data-generating parameters. The inference step estimates the posterior distribution of parameters for each simulated dataset using the same model as was used for simulation. For comparison, we compute Bayesian HPD intervals across all datasets for each parameter to assess whether we attain the targeted level of accuracy; if 95%-HPDs are used, for example, then coverage should be approximately 95% for each sampled parameter.

The model we validated was a three-region (a seven-range model), two-epoch time-heterogeneous GeoSSE model (TimeFIG). Our simulations assumed diversification began with one species at age $t = 2$, evolving under rates defined by the first epoch until age $t = 1$, with the process stopping at the present at age $t = 0$. We used rejection sampling to generate trees containing between 2 and 500 taxa. Base rates for $\rho_w$ and $\rho_b$ were independently drawn from LogNorm(0, 0.5), whereas $\rho_d$ and $\rho_e$ were independently drawn from LogNorm(-2, 0.5). Feature effect parameters ($\phi_p$ and $\sigma_p$ for the four processes $p \in \{d, e, w, b\}$) were independently sampled from Norm(0,1) that was truncated to [-1, 1]. We used one regional feature layer of each kind (four total) with artificial values that changed across two epochs (Table S6). We then independently inferred the posterior density for 100 simulated datasets under a TimeFIG model where all feature effect parameters were enabled and conditioning on the simulated tree as being true. We assessed the coverage of the base rate ($\rho_p$), quantitative feature effect ($\phi_p$), and categorical feature effect ($\sigma_p$) parameters associated with the four processes $p \in \{d, e, w, b\}$.

Estimated coverage was acceptably close to the target of 95% for all parameters (Figure S2). We reiterate that the purpose of this experiment was to validate the model implementation, and not to assess the statistical power to detect parameter effects (see Figs. S4, S5, and S6). That said, estimates for parameters associated with dispersal and within-region speciation were more precise than estimates for between-region. The simulated datasets contained little detectable information about extinction rates. Previous work also found weak extinction and between-region speciation feature effects were difficult to estimate precisely with time-homogenous FIG models, with more regions and larger trees leading to increased statistical power [10, 19].

#### Dating experiment

We simulated phylogenetic and biogeographic datasets to measure how well TimeFIG infers the crown age of a Hawaiian radiation. We sought island radiation datasets with small-to-medium sized and densely-sampled clades nested within older sparsely-sampled continental clades. In addition, dispersal between the continent and island must be rare to limit movement and create endemic radiations. Our initial efforts to simulate datasets resembling the properties of the *Kadua* dataset with standard techniques and brute force rejection sampling were not successful, as it required us to run hundreds of time-consuming simulations of large phylogenies to obtain only a few datasets that resembled the *Kadua* radiation. This motivated us to develop an alternative strategy that gave us finer control over simulation properties. Our simulation procedure had four steps: (1) simulating the timescale of key events, (2) simulating the island radiation, (3) adding the outgroup taxon, and (4) simulating sequence data.

First, to simulate the timescale of key events, we assumed that each radiation began with a single mainland lineage colonizing the archipelago at a random time under one of two scenarios. The first scenario assumed recent colonization into one of the modern islands, $t_{col} \sim \text{Unif}(1, 6)$. The second scenario assumed older colonization predating the modern islands, $t_{col} \sim \text{Unif}(6, 20)$. After sampling $t_{col}$, we then sampled a random crown age for the island radiation, $t_{crown} \sim \text{Unif}(1, t_{col})$, that was younger than the colonization time. Next, we simulated a random root age, $t_{root} \sim \text{Unif}(t_{col}, 32)$ for the continental clade containing the island radiation.

Second, we simulated the island radiation with an origin age, $t_{crown}$, under the TimeFIG model. We chose feature effect parameter values that allowed regional features to shape biogeographic rates in several important ways, including: distance decreased dispersal rates, long-distance dispersal into and out of the archipelago was rare, extinction rates increased in older islands, and so on. We used the dispersal rates determined by these feature effects and the paleogeographic features to sampled each ingroup's ancestral region, $s_{col}$, using the relative rate of dispersal from mainland region $Z$ into region $i$ at time $t_{col}$. We retain the time-calibrated phylogeny and biogeographic range data for the ingroup clade. For base rates, we set $\rho_d = 0.025$, $\rho_e = 0.5$, $\rho_w = 0.25$, and $\rho_b = 2$ to place our region-specific biogeographic rates on an absolute timescale that produced realistic biogeographic patterns. Feature effect parameters were set to: $\phi_w^{size} = 0.25$, $\phi_e^{size} = -1.0$, $\phi_w^{age} = -0.25$, $\phi_e^{age} = 0.5$, $\phi_d^{dist} = -1.0$, $\phi_b^{dist} = 2.0$, $\phi_d^{\Delta age} = -0.25$, $\phi_b^{\Delta age} = 0.0$, $\sigma_w^{high} = 0.5$, $\sigma_e^{high} = -2.0$, $\sigma_w^{grow} = 0.0$, $\sigma_e^{grow} = -0.5$, $\sigma_d^{dist} = -3.0$, $\sigma_b^{dist} = 2.0$, $\sigma_d^{\Delta age} = 0.5$, $\sigma_b^{\Delta age} = 0.0$. We used rejection sampling to simulate clades restricted to 25 to 75 extant taxa.

Third, we added the outgroup taxon that was sister to the island ingroup clade. To do this, we grafted two lineages on to the root node with age $t_{root}$: one daughter lineage attaches to the crown node for the island clade, and the other lineage is extended to the present as a singleton, representing an outgroup taxon. We associate the outgroup taxon with the outgroup region, Z. This gives us a single time-calibrated, ultrametric tree that contains all ingroup taxa plus one outgroup taxon.

Lastly, we simulated sequence evolution for 2,000 nucleotide sites under Jukes-Cantor substitution process with independent log-normally distributed branchwise clock rate variation. Longer sequences with more complex molecular substitutions are not necessary to show that TimeFIG can be used to time-calibrate phylogenies. The base clock rate was modeled as $\mu_{mol} \sim \text{Logunif}(5e-4, 5e-2)$, and the branch-specific clock rates were modeled as $\mu_j^{branch}$, $\mu_j^{branch} \sim \text{Lognorm}(ln(\mu_{mol}) - \frac{1}{2}\sigma_{mol}^2, 0.587405)$, such that their expected value was $\mu_{mol}$ and 95% of clock rate variation falls within one order of magnitude.

Under this procedure, we generated 100 datasets assuming a younger colonization time (1 to 6 Ma) and another 100 datasets assuming an older colonization time (6 to 20 Ma). We then analyzed each dataset twice: first under the full TimeFIG model (M1) and again under an uncalibrated model (M0) that assumes all feature effect parameters are disabled (=0) and that all islands always existed.

During inference, we used the following priors. The base rate $\rho_p$ for each process $p \in \{d, e, w, b\}$ was distributed by Exp(20). Feature effect parameters under M1 were modeled using reversible jump MCMC with probability 1/2 equal to zero and probability 1/2 of being distributed by Norm(0,1). We assumed taxon sampling for region Z was equal to $1/\tilde{E}(N_{t_{root}})$ where $\tilde{E}(N_{t_{root}}) = 2 \times \exp\left[t_{root}(r_w(Z, 1) - r_e(Z, 1))\right]$ is used to roughly approximate the expected number of species in region Z after time $t_{root}$. The root state frequencies required the root node to occupy region Z but allowed for the ancestor to simultaneously occupy one of any other island regions that existed at $t_{root}$. Priors for the molecular substitution process followed the simulating distributions described above.

After analyzing the simulated data for the young and old colonization scenarios with M0 and M1 using RevBayes, we computed posterior means and HPD95 credible intervals of island crown node age to compare the precision and accuracy of models M0 and M1, where M1 should consistently outperform M0. Because ourinference models both misspecify our fairly complicated data-generating model, we do not expect that 95% of HPD95s for M0 or M1 will contain the true ingroup age. We found that in general, the TimeFIG model used for M1 accurately estimated ingroup crown ages for the island radiation, whereas M0 consistently overestimated ingroup crown ages. Under the young colonization scenario, M1 estimated the correct age in 83% of cases, underestimated the age in 8.5% of cases, and overestimated the age in 8.5% cases, whereas M0 always overestimated the true age. For the old colonization scenario, M1 inferred the correct age in 80% of cases, underestimated the age in 16% of cases, and overestimate the age in 4% of cases. M0 inferred the correct age in 40%, particularly for true clade ages older than 10 Ma. Since M0 is uncalibrated, our interpretation is that this only occurs when the prior and posterior expectations happen to align by chance. In addition, the mean squared error and credible interval widths are much smaller for M1 than M0, indicating greater accuracy and precision for TimeFIG. Overall, these lend credibility ingroup clade age estimates we obtained in the empirical analysis of Hawaiian *Kadua*.

#### Parameter estimation experiment

We used simulations to assess how reliably TimeFIG estimates feature effect parameters under empirical conditions resembling the *Kadua* analysis. To do so, we simulated trees containing 20 to 50 terminal taxa, using the Hawaiian regions and regional features as defined for the empirical study. During simulation, we set the process base rates $\rho_d = 0.5$, $\rho_w = 0.05$, $\rho_e = 0.05$, and $\rho_b = 0.2$. We also set each of the 16 feature effect parameters to have positive or negative values as follows: $\phi_w^{size} = \phi_d^{\Delta age} = +0.5$; $\phi_e^{age}, \phi_b^{dist} = +1.0$; $\sigma_w^{high}, \sigma_b^{dist}, \sigma_e^{grow}, \sigma_d^{\Delta age} = +1.5$; $\phi_w^{age} = \phi_d^{dist} = -0.5$; $\phi_e^{age}, \phi_b^{\Delta age} = -1.0$; $\sigma_e^{high}, \sigma_d^{dist}, \sigma_w^{grow}, \sigma_b^{\Delta age} = -1.5$. The positive vs. negative signs for feature effects represent biogeographically plausible relationships between regions and hypotheses (e.g. distance decreases dispersal). For each parameter and dataset, we then randomly turned 'Off' each parameter (=0) with probability 0.5, meaning on average each dataset had 8 of 16 feature effects active. Next, we estimated the posterior distributions across all 200 datasets under the full model with the same priors as used for the dating experiment, but treating the phylogeny as known (fixed) to reduce run times.

We then used the posterior means and 80% HPDs for each replicate, and summarized performance by taking the mean-of-posterior-means and the mean of the squared errors posterior means relative to the scale of the 'On' value ($\frac{1}{N}\sum\left(\frac{x_{est}-x_{on}}{|x_{on}|}\right)^2$). We also classified estimates by HPD intervals in two ways. HPD intervals that include $x_{on}$ are labeled as having the correct value ('Value'), and intervals that exclude values with the *opposite* sign of $x_{on}$ are labeled as having the correct sign ('Sign'). We used this to report how frequently replicates recovered HPDs with the correct value and sign ('a'), only the correct sign ('b'), only the correct value ('c'), or the incorrect sign and value ('d'). Results were reported separately for datasets where each focal feature effect was truly 'On' versus 'Off', regardless of whether the remaining 15 feature effects were 'On' or 'Off'.

In general, estimates for datasets where the focal feature effect parameter was truly 'On' (Figure S4) had greater accuracy and higher proportions of HPDs with correct values and/or signs when compared to cases when the feature effects were 'Off' (Figure S5). Relative error terms under the 'On' condition were less than 0.65 for 10 parameters, but only for 1 parameter under the 'Off' condition. Similarly, more than 50% of intervals were correct in sign and/or value under the 'On' condition for 12 parameters, but only correct for 2 parameters under the 'Off' condition. This suggests that TimeFIG has the ability to estimate the correct value and/or sign of most feature effect parameters when the effect is truly present, but that the signal cannot always be detected.

#### Hypothesis testing experiment

Using the same simulated datasets and posteriors from the parameter estimation experiment, we computed Bayes factors supporting every hypothesis for every dataset ($200 \times 16 = 3200$ Bayes factors in total). For each simulated dataset, we knew whether each of the 16 hypotheses was true ('On') or false ('Off'), allowing us to compare the distributions of true positive and false positive Bayes factor scores, and compute false discovery scores, at different significance levels (Figure S6).

False discovery rates at the 'substantial' significance level (BF>3.2) for detecting a signed effect of a feature on a process were low (<10%) for six parameters ( $\phi_e^{size}$, $\phi_d^{dist}$, $\phi_b^{\Delta age}$, $\sigma_w^{high}$ $\sigma_w^{grow}$, $\sigma_b^{dist}$ ), medium (10−25%) for four parameters ( $\phi_b^{dist}$, $\phi_w^{age}$, $\phi_d^{\Delta age}$, $\sigma_d^{\Delta age}$ ), and high (25−40%) for three parameters ( $\phi_w^{size}$, $\phi_e^{age}$, $\sigma_d^{dist}$ ). FDRs for each of these 13 parameters fell to 0% at the 'strong' significance level (BF> 10), except for four parameters that had FDRs of ≤15%. Outside of those 13 parameters, the three remaining parameters ( $\sigma_e^{high}$, $\sigma_e^{grow}$, $\sigma_b^{\Delta age}$ ) were difficult to select or associate with FDRs regardless of the significance level. We emphasize that these are simulated for the Hawaiian system under a particular values of signed feature effects, depending on the parameter. These effects may be stronger or weaker than those influencing Hawaiian *Kadua*. However, the simulation results substantiate that small datasets similar to that for Hawaiian *Kadua* can generate detectable signal for biogeographic hypotheses testing, and that using Bayes factors to detect that signal is generally conservative.

#### TensorPhylo enhancements

For this project, we enhanced TensorPhylo to allow its phylogeny to be a random variable rather than a fixed value. This allows TensorPhylo to be used with RevBayes for tree inference, such as biogeographic dating with TimeFIG, fossil tip dating with an SSE model, or epidemiological modeling with a multitype birth-death process. As part of the enhancement, we added support to only recompute partial likelihoods impacted by local tree proposals to topology or divergence times (sometimes called "dirty flagging"). In principle, performance gains depend on tree size and tree balance, with large and balanced trees benefiting the most. For the small trees of with random topologies for roughly 20 taxa that we used for testing, dirty flagging produced a 2-4x speed-up. We validated correctness of the dirty flagging implementation by comparing the MCMC traces in RevBayes when using standard likelihood machinery versus the new TensorPhylo machinery. Under various conditions, both approaches produce the same sequence of posterior samples of phylogenetic trees when the random number generator is fixed to the same seed.

In addition, we added numerous safety checks to help the boost library's adaptive ordinary differential equation (ODE) solver avoid getting stuck on effectively impossible approximation problems. This problem sometimes arose with TimeFIG when MCMC would propose an extreme value for a feature effect parameter, which resulted in extreme degrees of rate asymmetry across states and time intervals. In addition, treating the tree as a random variable meant TensorPhylo would need to compute likelihoods for trees, also proposed by MCMC, with unusually short or long branches that could be difficult for the ODE solver to explain. In both case, the ODE solver could either hang indefinitely, unable to find a solution, or it could consume all system memory when attempting the solution, leading to software crashes. The new safety checks in TensorPhylo now predict when the ODE solver cannot find a solution in a reasonable amount of time, and sets the model likelihood to zero, causing the MCMC proposal to fail. In our testing environment, these incomputable model likelihoods are generally very rare (fewer than 1 in 1 million MCMC proposals), and forcing rejection for incomputable likelihoods had no discernible effect on results when experimenting with different levels of sensitivity to detect 'bad likelihoods'. These enhancements are available in commit 8887a4c of the main branch for TensorPhylo.

#### Supplementary Figures

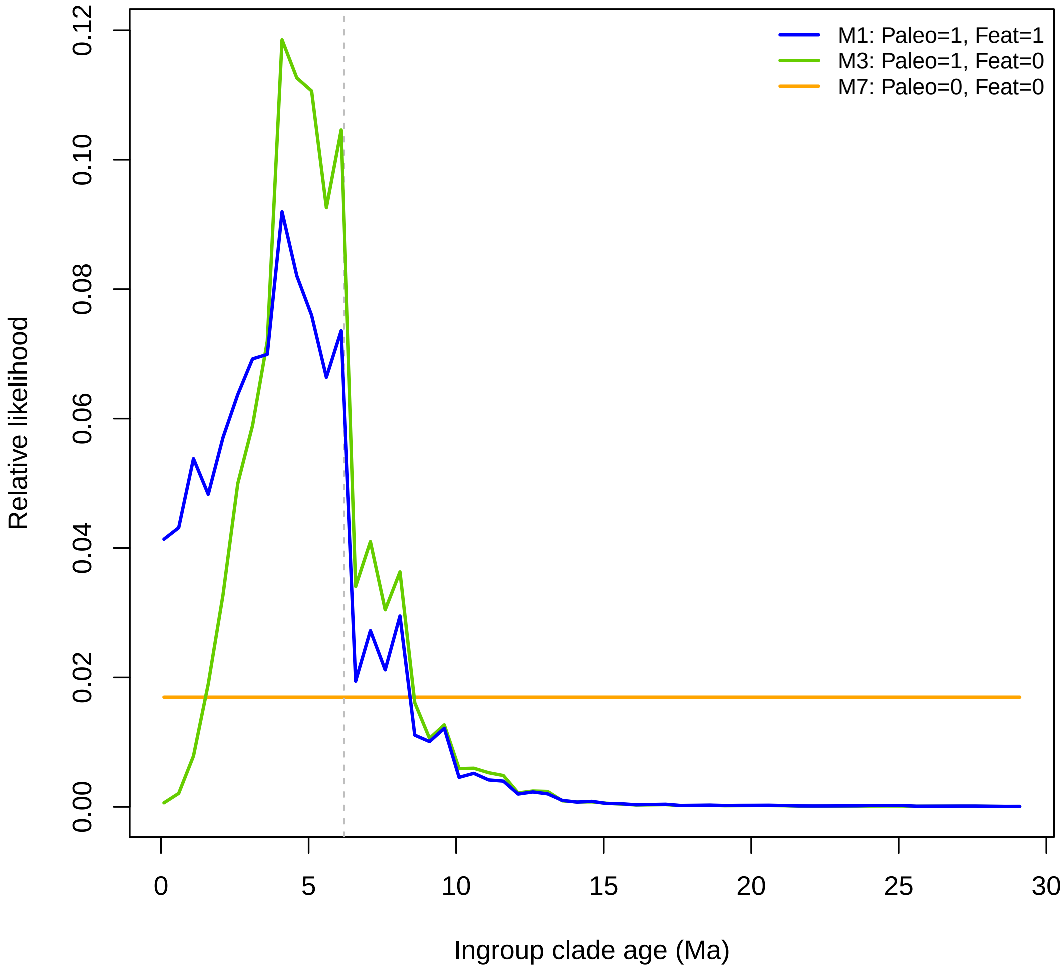


Figure S1: **Rate-time identifiability under TimeFIG model.** Demonstration of relative model likelihoods across different timescales for three model variants: a time-constant model that ignores paleogeographic island ages and regional features (gold; M7), a time-heterogeneous model that accounts for paleogeographic island ages but ignores regional features (green; M3), and a time-heterogeneous model that accounts for paleogeographic island ages and regional features (blue; M1). The vertical line represents the age estimate for Kauai, the oldest modern island. Details for how models were specified and relative likelihoods were computed are given in the Supplementary Text.

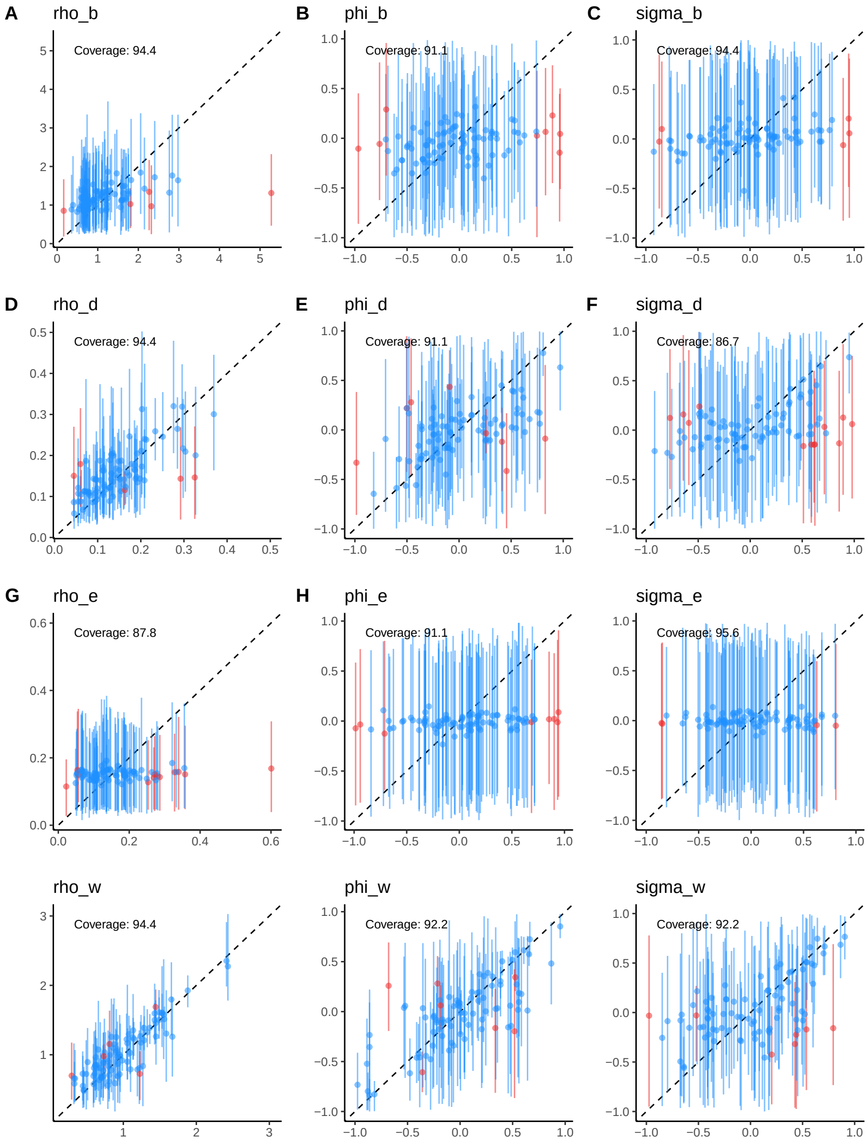


Figure S2: **Bayesian coverage estimates under TimeFIG model (3 regions, 2 timeslices).** Subplots in columns correspond to parameter types (base rates, quantitative effects, categorical effects). Subplots in rows correspond to process type (between-region speciation, dispersal, extinction, within-region speciation). Each subplot contains 100 replicates. Each point represents the true and posterior mean value for one simulated replicate. The dashed back line marks the cases where the true and estimated parameters match perfectly. Each vertical line represents the 95% credible interval for the highest posterior density (CI) for the replicate, and is marked blue if the CI includes the truth and red otherwise. These results were only generated to show that parameters estimates have acceptable coverage levels, near the 95% target level. Figures S4–S3) illustrate the ability of TimeFIG to test biogeographical hypotheses and estimate divergence times for the Hawaiian system (7 regions, 7 time periods).

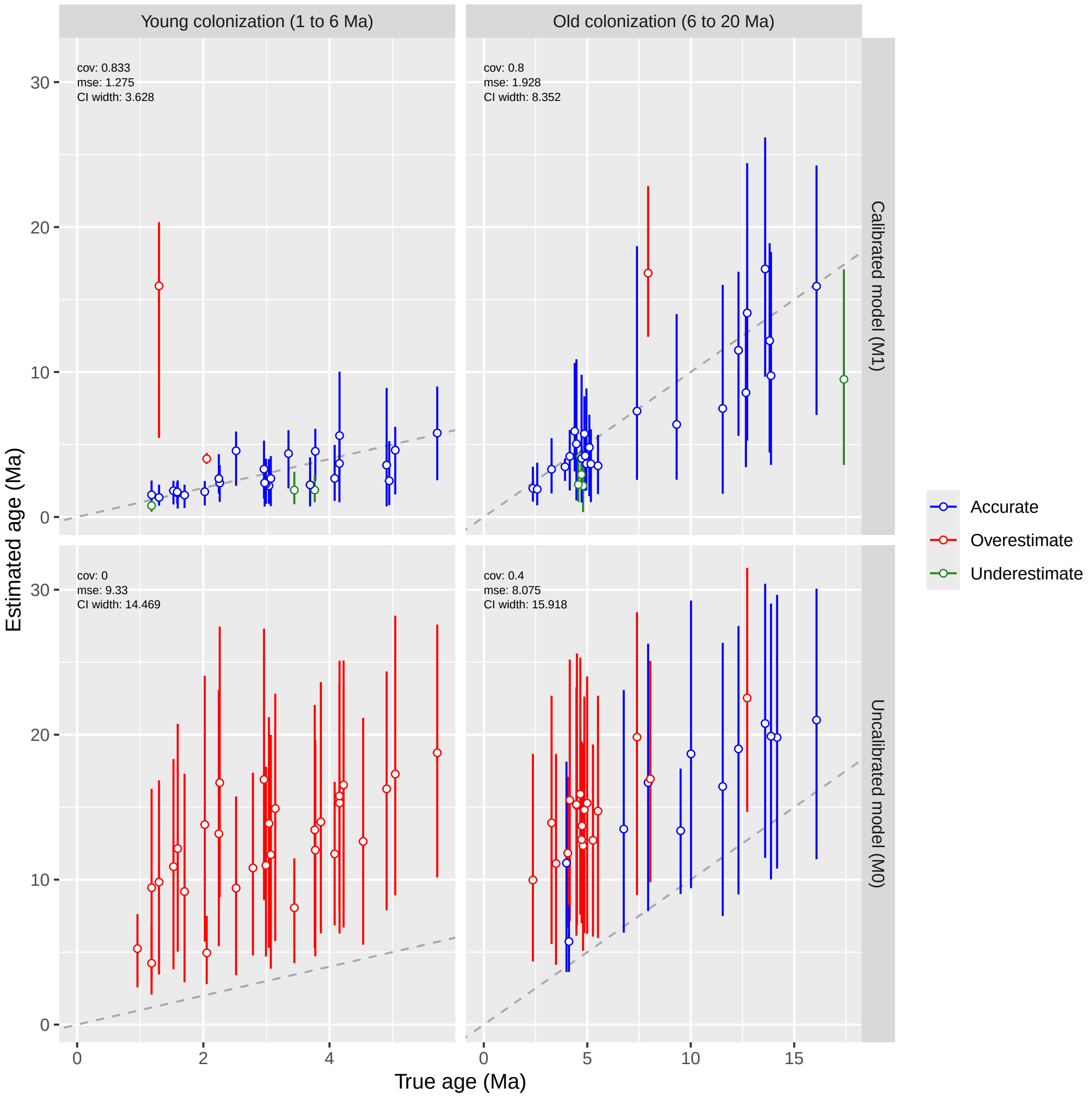


Figure S3: **Island clade age estimates under TimeFIG model.** Data were simulated to resemble the Hawaiian radiation for *Kadua* with young colonization times (1 to 6 Ma; left) and old colonization times (6 to 20 Ma; right), and then analyzed using TimeFIG (M1; top) and an uncalibrated model (M0; bottom); (Supplementary Text). Markers and bars represent the posterior means and HPD95 credible intervals. The gray line indicates when an estimate is equal to a true value. Each estimate was classified as accurate (blue) if the HPD95 contained the truth, an overestimate (red) if the HPD95 lower bound was greater than the truth, or an underestimate (green) if the HPD95 upper bound was less than the truth.

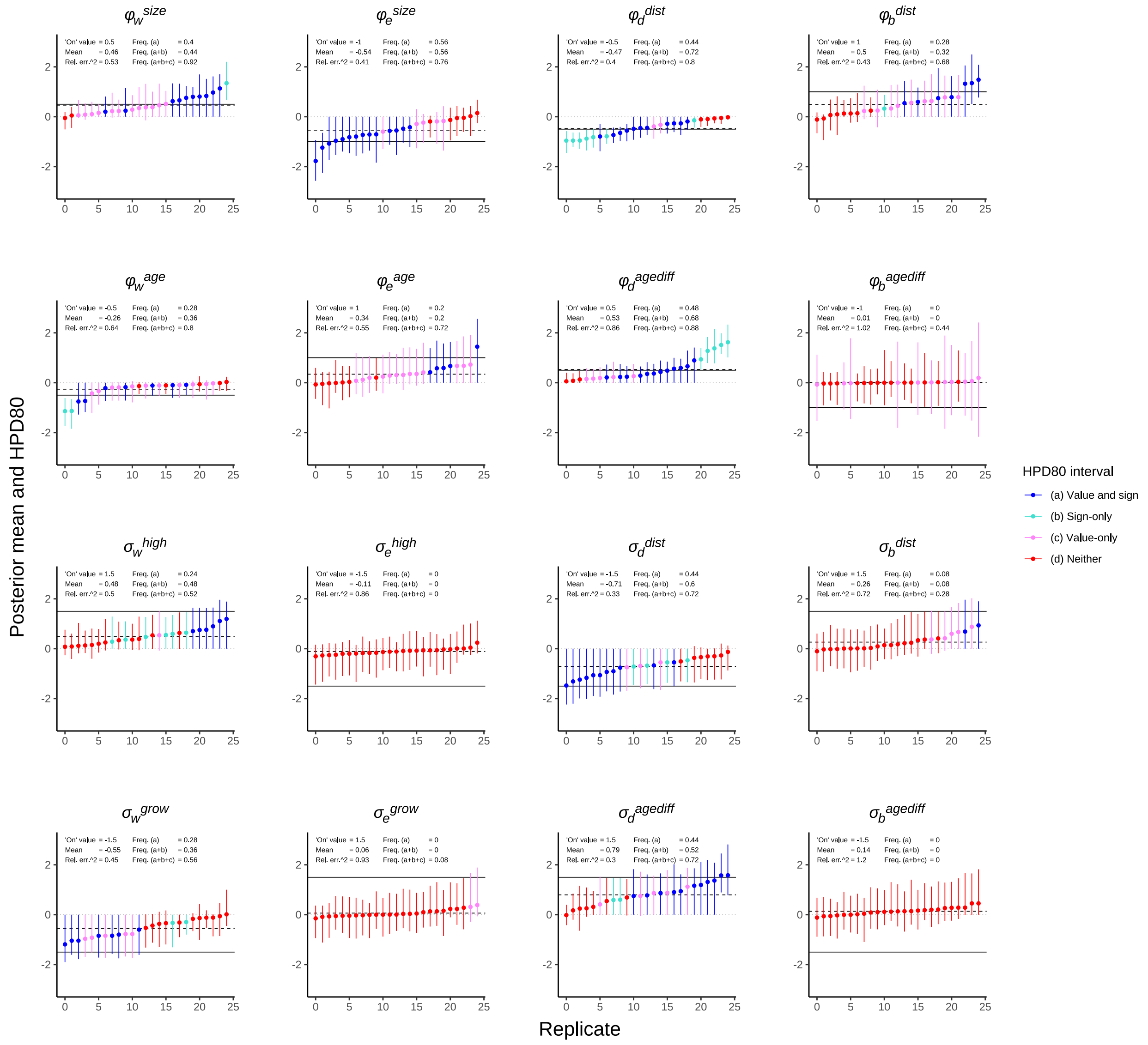


Figure S4: **Parameter estimation under TimeFIG for simulated Hawaiian radiations (focal effect is 'On').** Subplots report posterior means (points) and 80% highest posterior densities (bars) across 25 simulated datasets for the 16 feature effect parameters. The 16 subplots are arranged by the four process types (columns) and the eight regional features (rows). Within each subplot, each point represents a simulated dataset generated with that effect set to $\neq 0$ ('On') to follow its signed hypothesis, while each remaining effect is randomly 'On' or 'Off' (see text). Horizontal lines represent the feature effects values for the 'On' value (solid black), the mean-of-means (dashed black), and no effect at zero (gray) summarize overall performance. The HPD interval for each replicate is classified by whether it includes the 'On' value ('Value') and whether it excludes values with a sign opposite to the 'On' value ('Sign'), and then colored by scheme as Sign and Value (a: blue), Sign-only (b: turquoise), Value-only (c: pink), and Neither (d: red). The text reports the true value, the mean-of-means, the mean squared error relative to the scale of the 'On' value, and the frequency of HPD intervals (type a; type a or b; and type a, b, or c) across the full set of simulations, not just the 25 shown.

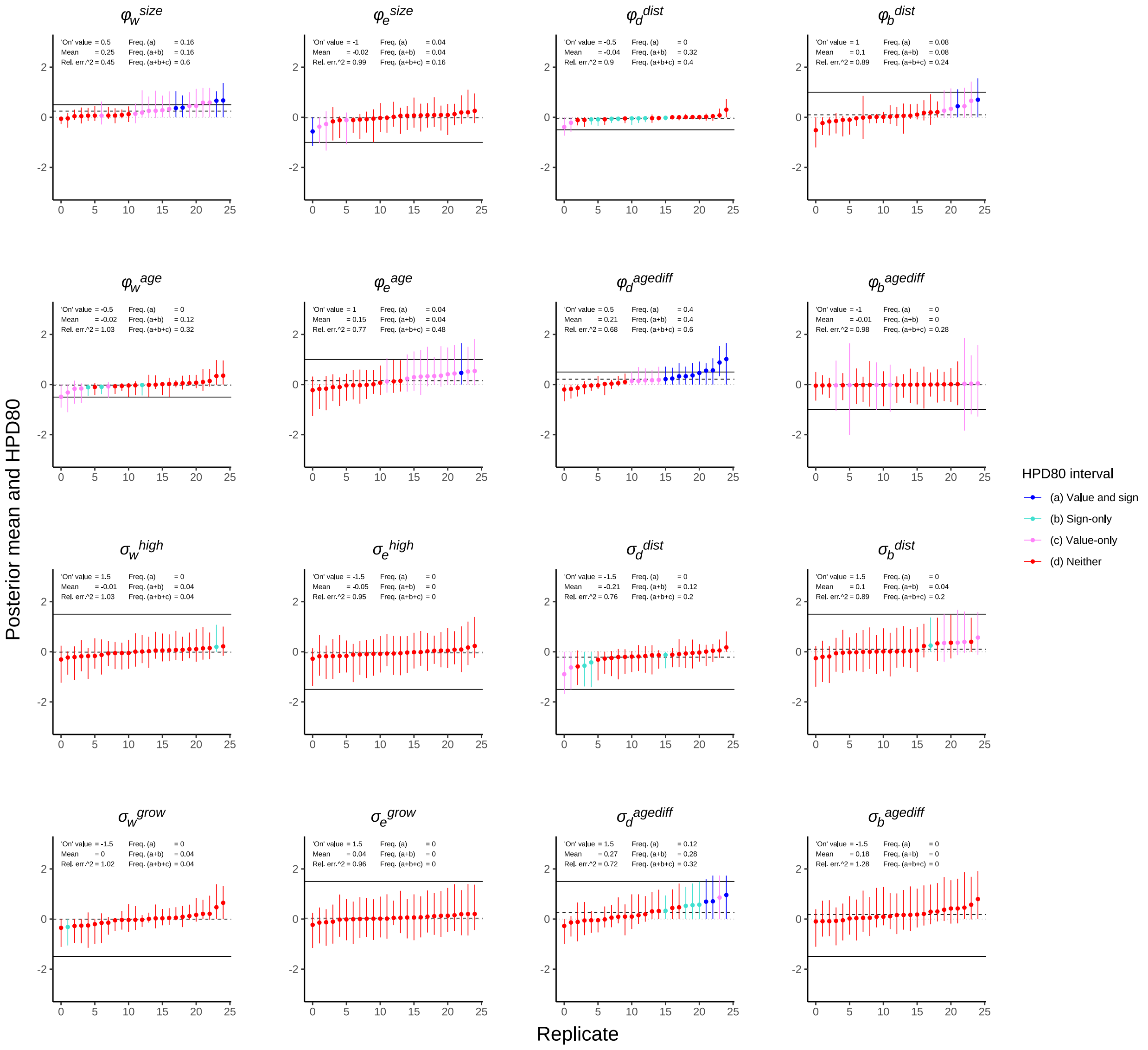

Figure S5: **Parameter estimation under TimeFIG for simulated Hawaiian radiations (focal effect is 'Off').** Subplots report posterior means (points) and 80% highest posterior densities (bars) across 25 simulated datasets for the 16 feature effect parameters. The 16 subplots are arranged by the four process types (columns) and the eight regional features (rows). Within each subplot, each point represents a simulated dataset generated with that effect set to $\neq 0$ ('On') to follow its signed hypothesis, while each remaining effect is randomly 'On' or 'Off' (see text). Horizontal lines represent the feature effects values for the 'On' value (solid black), the mean-of-means (dashed black), and no effect at zero (gray) summarize overall performance. The HPD interval for each replicate is classified by whether it includes the 'On' value ('Value') and whether it excludes values with a sign opposite to the 'On' value ('Sign'), and then colored by scheme as Sign and Value (a: blue), Sign-only (b: turquoise), Value-only (c: pink), and Neither (d: red). The text reports the true value, the mean-of-means, the mean squared error relative to the scale of the 'On' value, and the frequency of HPD intervals (type a; type a or b; and type a, b, or c) across the full set of simulations, not just the 25 shown.

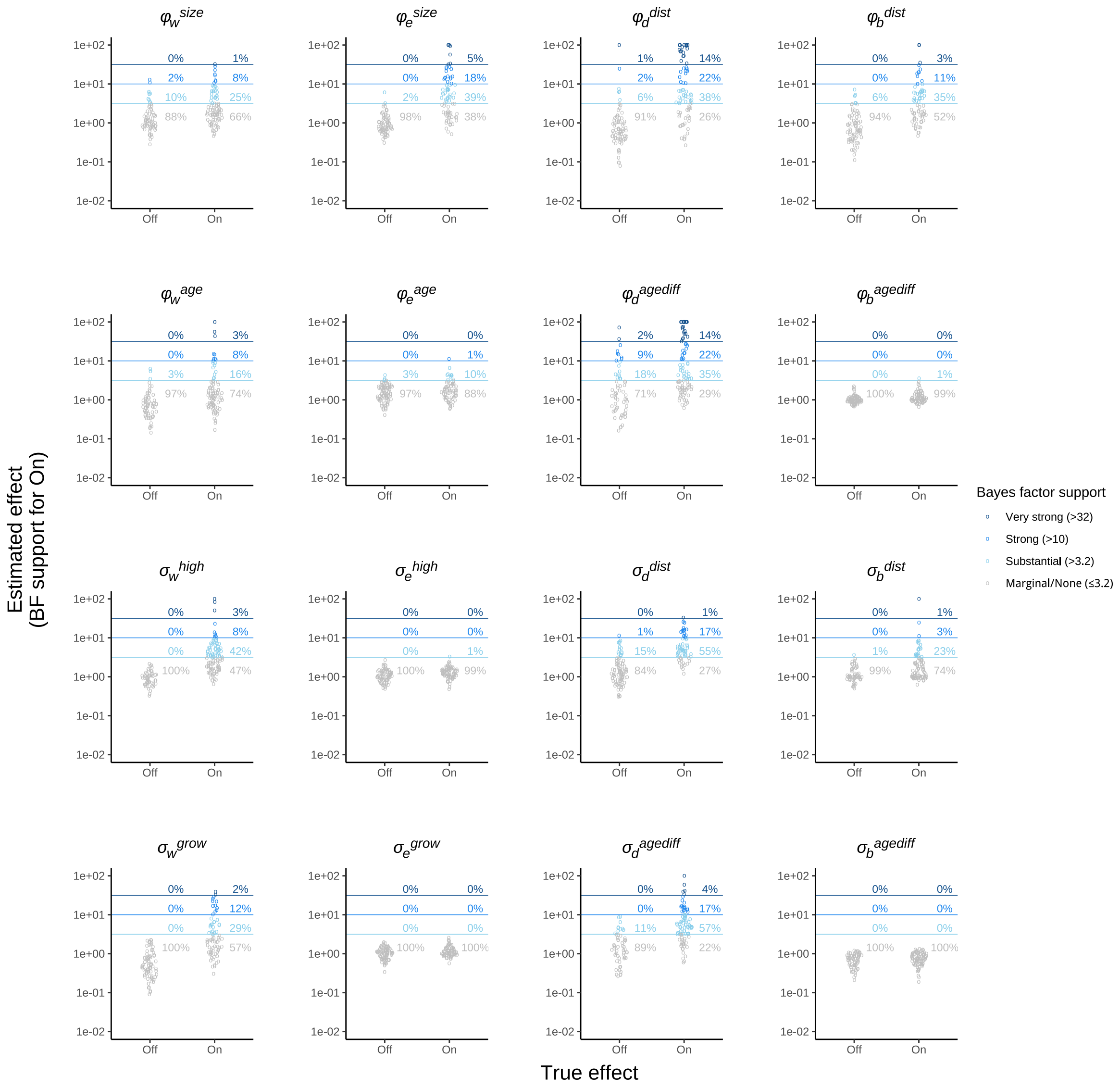


Figure S6: **Hypothesis testing under TimeFIG for simulated Hawaiian radiations.** Subplots report Bayes factor support (y-axis) for each feature effect parameter supporting a signed hypothesis. The 16 subplots are arranged by the four process types (columns) and the eight regional features (rows). Within each subplot, each point represents a simulated dataset generated while assuming each feature effect is randomly set to $= 0$ ('Off') or $\neq 0$ ('On') to follow its signed hypothesis (see text). Bayes factors are stratified (lines) into significance levels indicating little ($\leq 3.2$), substantial ($>3.2$, light blue), strong ($>10$, med. blue), or very strong ($>32$, dark blue) support for each signed hypothesis. Percentages report the proportion of inferences with significant support across datasets for which the feature effect was truly off vs. on during simulation. Sensitivity (true positive rate) is high when Bayes factor support is high among datasets simulated with the feature effect parameter set to on. False discovery rates are low when Bayes factor support is more common among datasets simulated with the feature effect parameter set to on, relative to those simulated with it set to off.

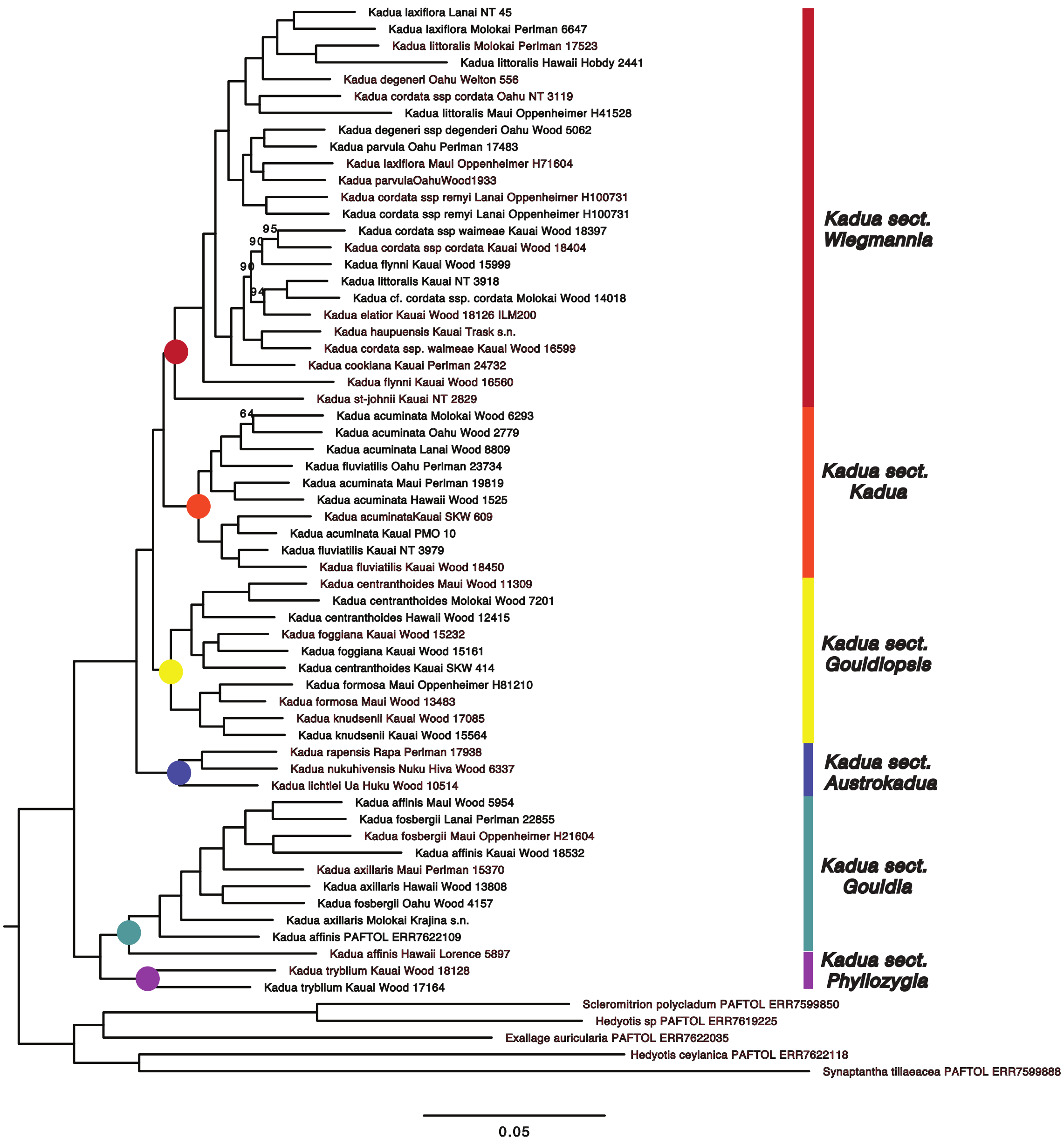


Figure S7: **Phylogenetic relationships of Hawaiian *Kadua* based on a concatenated data matrix of 343 low copy nuclear loci analyzed using Maximum likelihood in IQTREE 2**. Bootstrap support values less than 95 are shown at nodes.

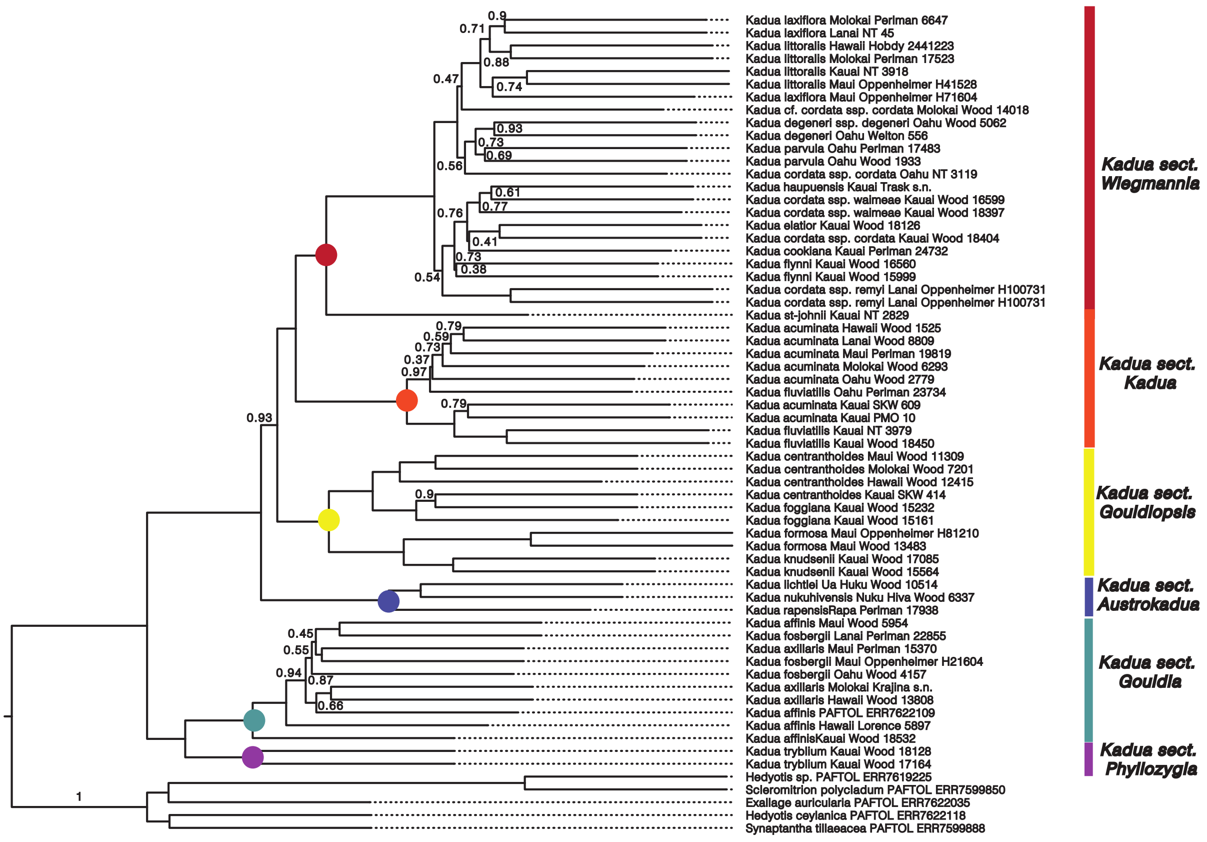


Figure S8: **Phylogenetic relationships of Hawaiian *Kadua* based on 343 low copy nuclear loci analyzed using Astral-III**. Local posterior probabilities less than 0.95 are shown at nodes.

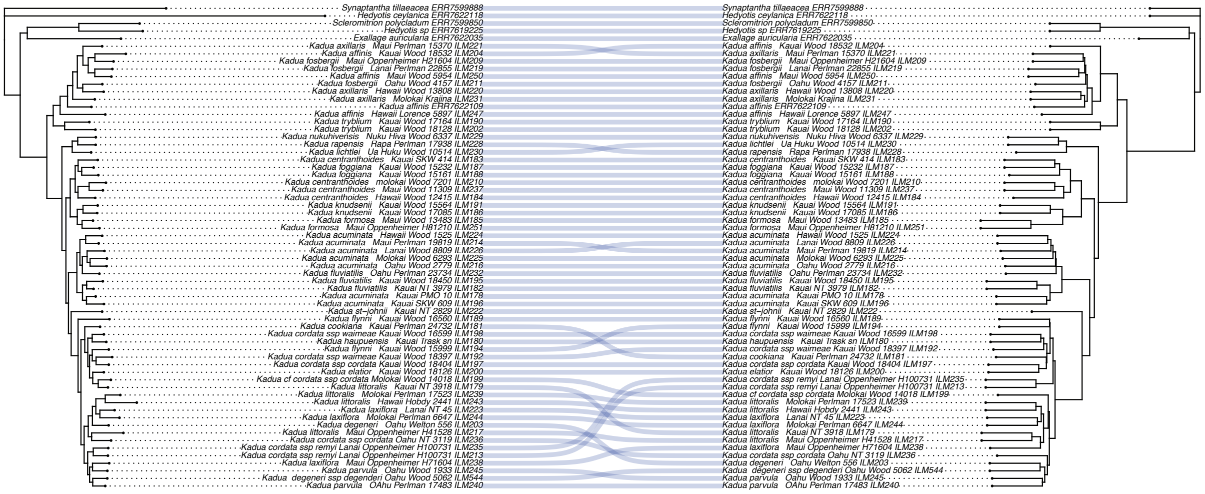

Figure S9: **Cophyloplot of Astral and Maximum Likelihood phylogenetic tree topologies of Hawaiian *Kadua* showing points of incongruence between the two topologies.**

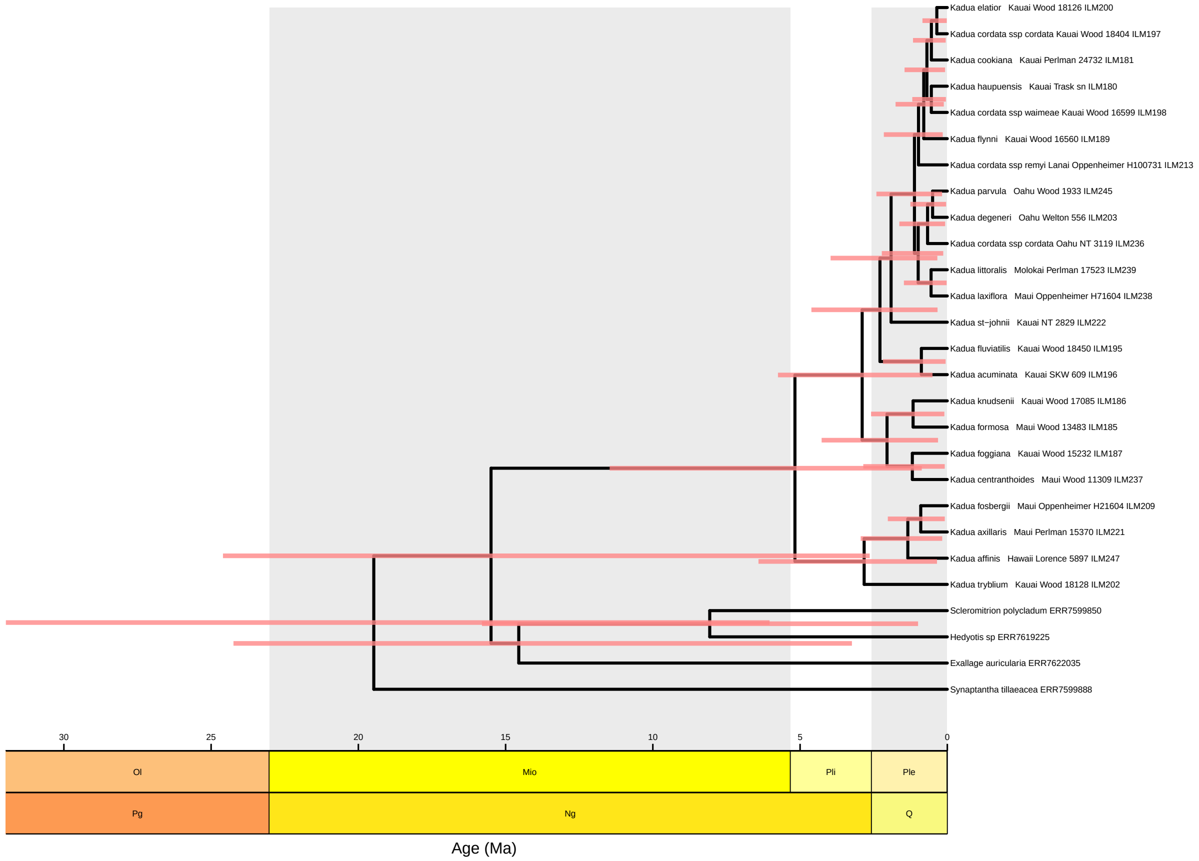


Figure S10: **Maximum clade credibility tree of Hawaiian *Kadua* for model M1.** Red bars represent 95% highest posterior density intervals for internal node ages.

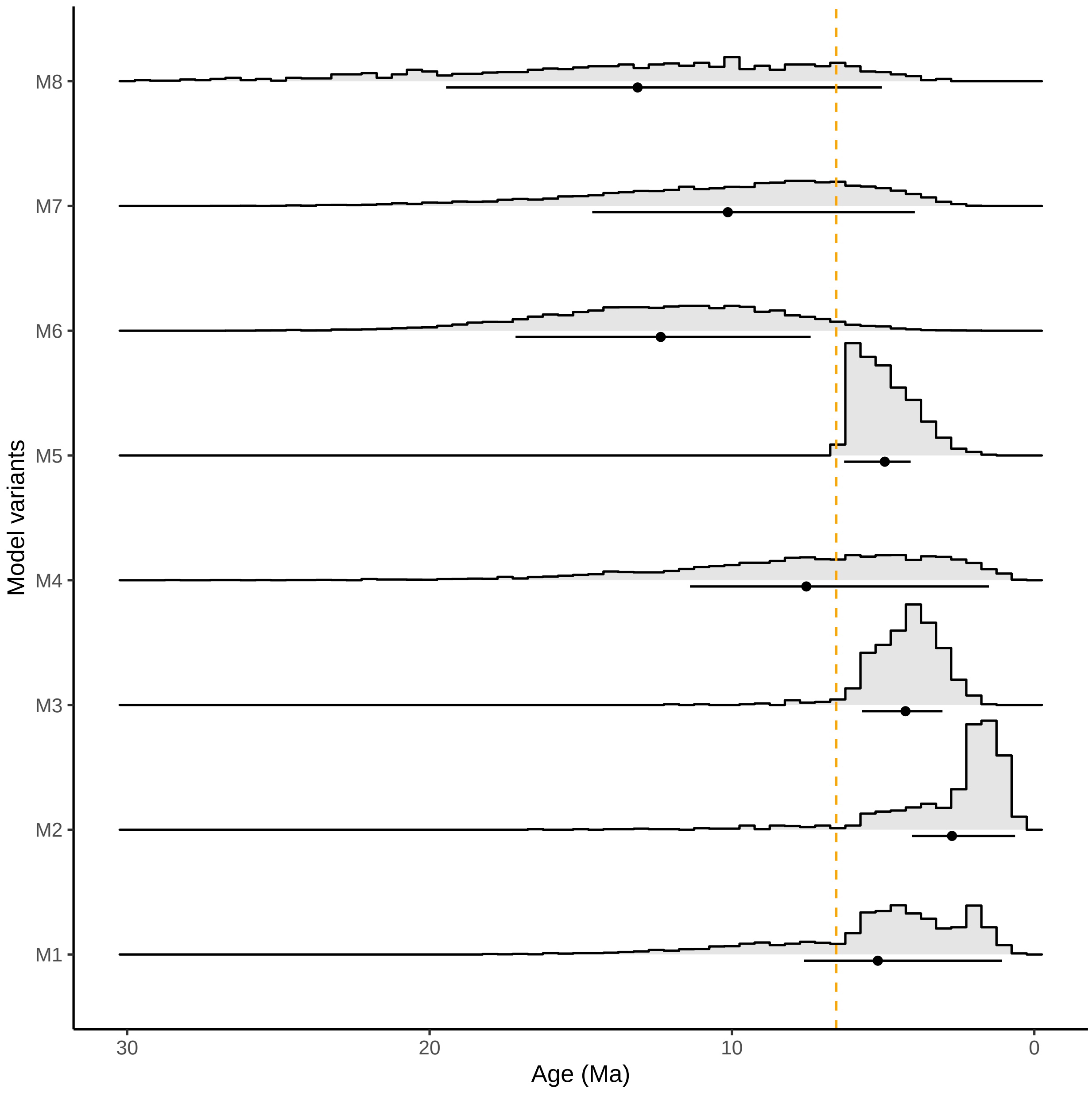


Figure S11: **Posterior mean crown ages of Hawaiian *Kadua* under the eight alternative model settings, M1–M8.** The point represents the posterior mean, and the bar represents the 95% highest posterior density credible interval. The yellow line corresponds 6.3 Ma, which is the maximum reported age of the Kaua'i + Ni'ihau region [11].

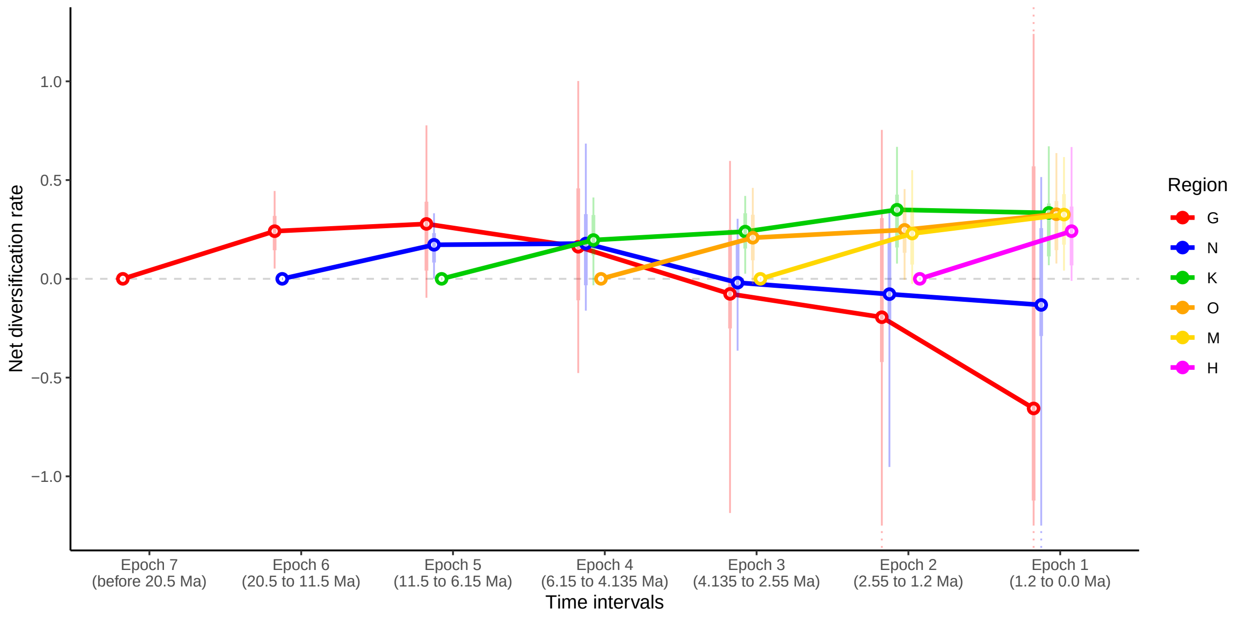


Figure S12: **Regional rates of net diversification over time.** Posterior densities of regional net diversification rates in region $i$ at time $t$ $(r_w(i,t) - r_e(i,t))$ from past (left) to present (right) at different stages of Hawaiian paleogeographic history. Rate variation among time intervals and regions is determined by feature effect parameter values interacting with paleogeographic changes in underlying regional features. Points represent posterior median rates for islands that existed during each time interval. The net diversification rate is assumed to be zero before the island exists. Solid lines trace how rates change between time intervals. Vertical lines give HPD50 (thick) and HPD80 (thin) intervals; dashed lines indicate HPD80 intervals that extend outside the scale of the figure. The non-Hawaiian region, Z, and the oldest epoch, when only Z existed, are not shown.

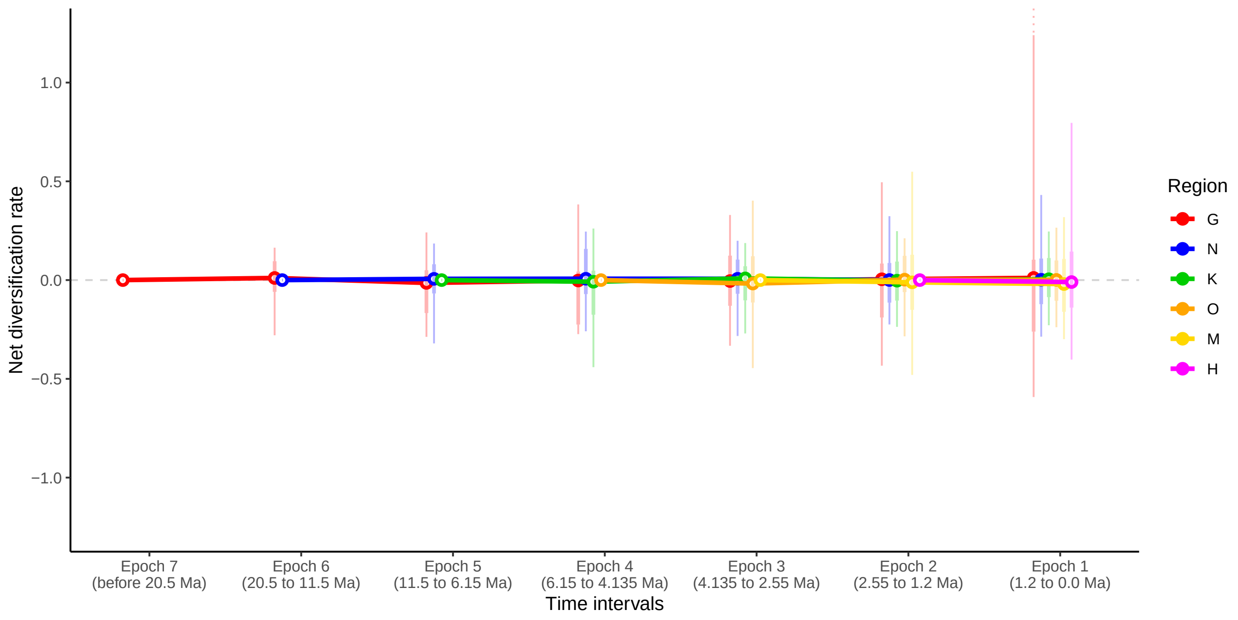


Figure S13: **Regional rates of net diversification over time, without phylogenetic or biogeographic data.** Prior densities of regional net diversification rates in region $i$ at time $t$ $(r_w(i,t) - r_e(i,t))$ from past (left) to present (right) at different stages of Hawaiian paleogeographic history. Rate variation among time intervals and regions is determined by feature effect parameter values interacting with paleogeographic changes in underlying regional features. Points represent posterior median rates for islands that existed during each time interval. The net diversification rate is assumed to be zero before the island exists. Solid lines trace how rates change between time intervals. Vertical lines give HPD50 (thick) and HPD80 (thin) intervals; dashed lines indicate HPD80 intervals that extend outside the scale of the figure. The non-Hawaiian region, Z, and the oldest epoch, when only Z existed, are not shown.

#### Supplementary Tables

Table S1: **Properties of molecular dataset.** Mean, median, and standard deviations of different properties for the 344 low-copy nuclear loci (supercontigs) used to infer the gene trees and species tree of *Kadua*. Gene and gene tree properties are based on *genesortR* method [15].

| Property | Variable(s) | Mean | Median | Std. dev. |
|---|---|---|---|---|
| Root-tip (variance) | Gene trees | 0.004 | 0.003 | 0.003 |
| Missing data | Alignments | 0.027 | 0.026 | 0.011 |
| Alignment length | Alignments | 1105.914 | 1002 | 682.157 |
| Occupancy | Alignments | 0.992 | 1 | 0.015 |
| Variable sites (proportion) | Alignments | 0.429 | 0.435 | 0.095 |
| Saturation | Alignments & gene trees | 0.313 | 0.318 | 0.101 |
| Evolutionary rate | Gene trees | 0.014 | 0.015 | 0.005 |
| Tree length | Gene trees | 0.794 | 0.803 | 0.297 |
| Treeness | Gene trees | 0.338 | 0.323 | 0.132 |
| Patristic dist. (mean) | Gene trees | 0.075 | 0.076 | 0.031 |
| Bootstrap support (mean) | Gene trees | 70.897 | 71.604 | 7.699 |
| RF similarity | Gene trees vs. species tree | 0.113 | 0.094 | 0.069 |

Table S2: **Island regions and ages**. Epochs are ordered by increasing starting ages, from 1 to 7, where the age column represents the midpoint between the maximum and minimum estimated ages from [11].

| Region | Symbol | Islands | Epoch | Age (Ma) |
|---|---|---|---|---|
| Hawai'i | H | | 1 | 1.2 |
| Maui Nui | M | Maui, Moloka'i, Lana'i, and Kaho'olawe | 2 | 2.55 |
| O'ahu | O | | 3 | 4.135 |
| Kaua'i | K | Ni'ihau and Lehua | 4 | 6.15 |
| Necker | N | Necker through Nihoa and Kalau | 5 | 10.6 |
| Gardner | G | Maro through Lisianski | 6 | 19.7 |
| – | – | (no habitable islands) | 7 | 32 |

Table S3: **Biogeographic hypotheses and paleogeographic features.** The four hypotheses (H1–H4) are described in the main text. Features can be within regions (W) or shared between regions (B), and have categorical (C) or quantitative (Q) values. Citations: C96 [2]; LM17 [11]; PC02 [17].

| Hypothesis | Within or between? | Categorical or quantitative | Data | Citations |
|---|---|---|---|---|
| Size-extinction (H1) | W | Q | Elevation (m) | PC02 |
| Distance-dispersal (H2) | B | C | Long-distance dispersal? | – |
| Distance-dispersal (H2) | B | Q | Geographic distance (km) | Google Earth |
| Progression rule (H3) | B | C | Into younger island? | – |
| Progression rule (H3) | B | Q | Island age differences (Ma) | C96, LM17 |
| Dynamic equilibrium (H4) | W | C | Is high island? | C96, LM17 |
| Dynamic equilibrium (H4) | W | C | In growth phase? | C96, LM17 |
| Dynamic equilibrium (H4) | W | Q | Island age (Ma) | C96, LM17 |

Table S4: **Biogeographic hypotheses and TimeFIG parameter values.** Summary of TimeFIG feature effect parameter values that would support the four key biogeographic hypotheses tested in the main text.

| Hypothesis | Parameters | Interpretation |
|---|---|---|
| Size-diversification (H1) | $\phi_w^{size} > 0$ | Speciation faster in large islands |
| | $\phi_e^{size} < 0$ | Extinction faster in small islands |
| Distance-dispersal (H2) | $\phi_d^{dist} < 0$ | Dispersal faster over short distances |
| | $\sigma_d^{dist} < 0$ | Dispersal faster among islands |
| Progression rule (H3) | $\phi_d^{\Delta age} > 0$ | Dispersal faster into younger islands (by amt.) |
| | $\sigma_d^{\Delta age} > 0$ | Dispersal faster into younger islands (any amt.) |
| Dynamic equilibrium (H4) | $\sigma_w^{high} > 0$ | Speciation faster in high islands |
| | $\sigma_w^{grow} < 0$ | Speciation faster in eroding/subsiding islands |
| | $\sigma_e^{high} < 0$ | Extinction faster in low islands |
| | $\sigma_e^{grow} > 0$ | Extinction faster in growing islands |

Table S5: **Model variant properties.** Summary of the model properties that are turned on (check mark) or off (no check mark) corresponding to variants M1 to M8. Supplementary Text describes these variants in greater detail.

| Model | Use data? | Use features? | Use paleogeo.? | Use 2° calibration? | Use biogeo. calibration? | Description |
|---|---|---|---|---|---|---|
| M1 | ✓ | ✓ | ✓ | ✓ | | Full process-dating analysis |
| M2 | ✓ | ✓ | ✓ | | | M1 + no secondary calibration |
| M3 | ✓ | | ✓ | | | M2 + no regional features |
| M4 | ✓ | ✓ | | | | M2 + no paleogeography |
| M5 | ✓ | | | | ✓ | Biogeographic calibration |
| M6 | ✓ | | | ✓ | | Secondary calibration |
| M7 | ✓ | | | | | Uncalibrated |
| M8 | | | | | | Uninformed (≈ under prior) |

Table S6: **Regional features for coverage analysis.** Values of regional features for three regions across two epochs to assess Bayesian coverage levels for simulated data. One layer was used for each of the four combinations of quantitative or categorical and within-region or between-region features ($q_w, c_w, q_b, c_b$).

| Feature | | Epoch 1 | | | Epoch 2 | | |
|---|---|---|---|---|---|---|---|
| | | A | B | C | A | B | C |
| $q_w$ | – | 5 | 10 | 35 | 25 | 10 | 5 |
| $c_w$ | – | 1 | 0 | 0 | 0 | 1 | 0 |
| $q_b$ | A | – | 10 | 30 | – | 50 | 30 |
| | B | 20 | – | 40 | 60 | – | 20 |
| | C | 60 | 80 | – | 80 | 10 | – |
| $c_b$ | A | – | 0 | 1 | – | 1 | 0 |
| | B | 0 | – | 1 | 1 | – | 0 |
| | C | 1 | 1 | – | 0 | 0 | – |

Table S7: **Parameter estimation experiment using simulated data.** Rows summarize parameter estimates for each focal parameter and the hypothesized value. Results are divided into those where the focal parameter equaled the hypothesized value ('On') or equaled zero ('Off') during simulation. Columns report the mean of posterior means, the mean squared error relative to the scale of the hypothesized value, and the frequencies of 80% highest posterior density intervals (HPDs) that include the hypothesized value (V1: yes, V0: no) and that exclude values with the opposite sign of the hypothesized value (S1: yes, S0: yes). See Supplement and Figures S4 and S5 additional information.

| | | Parameter 'On' | | | | | | Parameter 'Off' | | | | | |
|---|---|---|---|---|---|---|---|---|---|---|---|---|---|
| Param. | Hyp. value | Mean | Rel. err. | $f_{V1S1}$ | $f_{V0S1}$ | $f_{V1S0}$ | $f_{V0S0}$ | Mean | Rel. err. | $f_{V1S1}$ | $f_{V0S1}$ | $f_{V1S0}$ | $f_{V0S0}$ |
| $\phi_w^{size}$ | 0.5 | 0.36 | 0.46 | 0.4 | 0.42 | 0.01 | 0.16 | 0.18 | 0.57 | 0.12 | 0.43 | 0.08 | 0.37 |
| $\phi_e^{size}$ | -1.0 | -0.55 | 0.39 | 0.55 | 0.18 | 0.04 | 0.23 | -0.03 | 0.96 | 0.04 | 0.14 | 0.01 | 0.81 |
| $\phi_d^{dist}$ | -0.5 | -0.41 | 0.37 | 0.54 | 0.06 | 0.23 | 0.17 | -0.07 | 0.9 | 0.06 | 0.06 | 0.25 | 0.64 |
| $\phi_b^{dist}$ | 1.0 | 0.51 | 0.43 | 0.26 | 0.37 | 0.03 | 0.34 | 0.05 | 0.97 | 0.05 | 0.13 | 0.01 | 0.81 |
| $\phi_w^{age}$ | -0.5 | -0.2 | 0.69 | 0.28 | 0.33 | 0.05 | 0.34 | -0.01 | 1.13 | 0.06 | 0.19 | 0.05 | 0.7 |
| $\phi_e^{age}$ | 1.0 | 0.28 | 0.6 | 0.12 | 0.55 | 0 | 0.32 | 0.18 | 0.7 | 0.06 | 0.47 | 0 | 0.47 |
| $\phi_d^{\Delta age}$ | 0.5 | 0.57 | 0.79 | 0.39 | 0.25 | 0.28 | 0.08 | 0.23 | 0.7 | 0.39 | 0.13 | 0.05 | 0.43 |
| $\phi_b^{\Delta age}$ | -1 | -0.01 | 0.98 | 0 | 0.42 | 0 | 0.58 | -0.01 | 0.98 | 0 | 0.23 | 0 | 0.77 |
| $\sigma_w^{high}$ | 1.5 | 0.49 | 0.5 | 0.2 | 0.04 | 0.22 | 0.54 | 0.02 | 0.99 | 0 | 0 | 0.02 | 0.98 |
| $\sigma_e^{high}$ | -1.5 | -0.13 | 0.84 | 0.01 | 0 | 0 | 0.99 | -0.06 | 0.93 | 0 | 0 | 0.01 | 0.99 |
| $\sigma_d^{dist}$ | -1.5 | -0.76 | 0.3 | 0.41 | 0.18 | 0.11 | 0.3 | -0.22 | 0.76 | 0.03 | 0.04 | 0.1 | 0.82 |
| $\sigma_b^{dist}$ | 1.5 | 0.32 | 0.7 | 0.11 | 0.24 | 0.01 | 0.64 | 0.08 | 0.91 | 0 | 0.09 | 0.02 | 0.9 |
| $\sigma_w^{grow}$ | -1.5 | -0.45 | 0.57 | 0.21 | 0.09 | 0.08 | 0.61 | 0.09 | 1.15 | 0 | 0 | 0.05 | 0.95 |
| $\sigma_e^{grow}$ | 1.5 | 0.07 | 0.92 | 0 | 0.1 | 0 | 0.9 | 0.01 | 0.99 | 0 | 0.01 | 0 | 0.99 |
| $\sigma_d^{\Delta age}$ | 1.5 | 0.74 | 0.34 | 0.43 | 0.16 | 0.13 | 0.29 | 0.21 | 0.78 | 0.07 | 0.04 | 0.11 | 0.79 |
| $\sigma_b^{\Delta age}$ | -1.5 | 0.13 | 1.19 | 0 | 0 | 0 | 1 | 0.18 | 1.28 | 0 | 0 | 0 | 1 |

Table S8: **Bayes factor experiment using simulated data.** Rows summarize numbers of true positives ($N_{TP}$) and false positives ($N_{FP}$), percents of true positives ($P_{TP}$) and false positives ($P_{FP}$), and false discovery rates ($FDR$). Columns report results for Bayes factors indicating substantial support (>3.2), strong support (>10), or very strong support (>32). See Supplement and Figure S6 additional information.

| | | Substantial BF>3.2 | | | | | Strong BF>10 | | | | | Very strong BF>32 | | | | |
|---|---|---|---|---|---|---|---|---|---|---|---|---|---|---|---|---|
| Parameter | Hyp. value | $N_{TP}$ | $N_{FP}$ | $P_{TP}$ | $P_{FP}$ | $FDR$ | $N_{TP}$ | $N_{FP}$ | $P_{TP}$ | $P_{FP}$ | $FDR$ | $N_{TP}$ | $N_{FP}$ | $P_{TP}$ | $P_{FP}$ | $FDR$ |
| $\phi_w^{size}$ | 0.5 | 26 | 10 | 30.6 | 15.4 | 27.8 | 8 | 2 | 9.4 | 3.1 | 20 | 1 | 0 | 1.2 | 0 | 0 |
| $\phi_e^{size}$ | -1 | 44 | 2 | 56.4 | 2.8 | 4.3 | 20 | 0 | 25.6 | 0 | 0 | 6 | 0 | 7.7 | 0 | 0 |
| $\phi_d^{dist}$ | -0.5 | 51 | 6 | 65.4 | 8.3 | 10.5 | 30 | 2 | 38.5 | 2.8 | 6.2 | 19 | 1 | 24.4 | 1.4 | 5 |
| $\phi_b^{dist}$ | 1 | 35 | 6 | 47.9 | 7.8 | 14.6 | 11 | 0 | 15.1 | 0 | 0 | 3 | 0 | 4.1 | 0 | 0 |
| $\phi_w^{age}$ | -0.5 | 17 | 3 | 19.5 | 4.8 | 15 | 9 | 0 | 10.3 | 0 | 0 | 3 | 0 | 3.4 | 0 | 0 |
| $\phi_e^{age}$ | 1 | 10 | 3 | 15.4 | 3.5 | 23.1 | 1 | 0 | 1.5 | 0 | 0 | 0 | 0 | 0 | 0 | 0 |
| $\phi_d^{\Delta age}$ | 0.5 | 51 | 18 | 57.3 | 29.5 | 26.1 | 32 | 9 | 36 | 14.8 | 22 | 21 | 2 | 23.6 | 3.3 | 8.7 |
| $\phi_b^{\Delta age}$ | -1 | 1 | 0 | 1.4 | 0 | 0 | 0 | 0 | 0 | 0 | 0 | 0 | 0 | 0 | 0 | 0 |
| $\sigma_w^{high}$ | 1.5 | 47 | 0 | 47.5 | 0 | 0 | 9 | 0 | 9.1 | 0 | 0 | 3 | 0 | 3 | 0 | 0 |
| $\sigma_e^{high}$ | -1.5 | 1 | 0 | 1.3 | 0 | 0 | 0 | 0 | 0 | 0 | 0 | 0 | 0 | 0 | 0 | 0 |
| $\sigma_d^{dist}$ | -1.5 | 47 | 15 | 77 | 16.9 | 24.2 | 15 | 1 | 24.6 | 1.1 | 6.2 | 1 | 0 | 1.6 | 0 | 0 |
| $\sigma_b^{dist}$ | 1.5 | 23 | 1 | 25 | 1.7 | 4.2 | 3 | 0 | 3.3 | 0 | 0 | 1 | 0 | 1.1 | 0 | 0 |
| $\sigma_w^{grow}$ | -1.5 | 30 | 0 | 40 | 0 | 0 | 13 | 0 | 17.3 | 0 | 0 | 2 | 0 | 2.7 | 0 | 0 |
| $\sigma_e^{grow}$ | 1.5 | 0 | 0 | 0 | 0 | 0 | 0 | 0 | 0 | 0 | 0 | 0 | 0 | 0 | 0 | 0 |
| $\sigma_d^{\Delta age}$ | 1.5 | 69 | 10 | 73.4 | 17.9 | 12.7 | 20 | 0 | 21.3 | 0 | 0 | 5 | 0 | 5.3 | 0 | 0 |
| $\sigma_b^{\Delta age}$ | -1.5 | 0 | 0 | 0 | 0 | 0 | 0 | 0 | 0 | 0 | 0 | 0 | 0 | 0 | 0 | 0 |

Table S9: **Posterior clade age estimates for Hawaiian *Kadua* for model M1**. Columns represent posterior means and the lower and upper bounds of the 80% and 95% highest posterior density credible intervals.

| | | HPD80 | | HPD95 | |
|---|---|---|---|---|---|
| Clade | Mean | Lower | Upper | Lower | Upper |
| Hawaiian *Kadua* | 5.17 | 1.07 | 7.62 | 0.86 | 11.47 |
| Sect. *Weigmannia* | 2.89 | 0.54 | 2.79 | 0.31 | 3.68 |
| Sect. *Kadua* | 0.88 | 0.13 | 1.31 | 0.06 | 2.04 |
| Sect. *Gouldiopsis* | 2.05 | 0.52 | 2.94 | 0.36 | 4.02 |
| Sects. *Gouldia* + *Phyllozygia* | 2.82 | 0.72 | 4.11 | 0.36 | 5.99 |
| *Scleromitrion* + *Hedyotis* | 8.07 | 2.19 | 12.62 | 0.95 | 15.45 |
| *Exallage* + *Hedyotis* | 14.55 | 7.83 | 22.75 | 3.31 | 23.88 |
| *Kadua* + *Hedyotis* | 15.49 | 8.46 | 23.85 | 3.37 | 24.83 |
| *Kadua* + *Synaptantha* | 19.48 | 12.68 | 31.90 | 6.42 | 32.00 |

Table S10: **Posterior parameter estimates for M1**. Columns represent posterior means, medians, and lower and upper bounds for the 80% and 95% highest posterior density credible intervals. Parameter values include samples where feature effects are exactly equal to zero as part of the reversible-jump MCMC analysis.

| | | | HPD80 | | HPD95 | |
|---|---|---|---|---|---|---|
| Variable | Mean | Median | Lower | Upper | Lower | Upper |
| $\rho_w$ | 1.69e-01 | 1.49e-01 | 3.74e-02 | 2.51e-01 | 3.19e-02 | 3.61e-01 |
| $\rho_e$ | 9.48e-02 | 7.75e-02 | 4.72e-04 | 1.50e-01 | 1.30e-05 | 2.36e-01 |
| $\rho_d$ | 3.07e-02 | 2.41e-02 | 3.65e-03 | 4.61e-02 | 1.92e-03 | 7.78e-02 |
| $\rho_b$ | 8.53e-02 | 6.53e-02 | 3.64e-04 | 1.34e-01 | 1.11e-05 | 2.31e-01 |
| $\phi_w^{size}$ | 2.02e-01 | 0.00e+00 | -6.60e-02 | 9.45e-01 | -6.23e-01 | 1.64e+00 |
| $\phi_w^{age}$ | 1.92e-01 | 0.00e+00 | -6.74e-02 | 7.56e-01 | -4.46e-01 | 1.43e+00 |
| $\phi_e^{size}$ | -7.62e-01 | -5.66e-01 | -2.33e+00 | 1.52e-02 | -2.97e+00 | 9.76e-01 |
| $\phi_e^{age}$ | 2.24e-01 | 0.00e+00 | -5.11e-01 | 1.31e+00 | -1.53e+00 | 1.94e+00 |
| $\phi_d^{dist}$ | -1.38e+00 | -1.39e+00 | -2.04e+00 | -8.57e-01 | -2.42e+00 | -3.95e-01 |
| $\phi_d^{\Delta age}$ | 4.98e-01 | 2.35e-01 | 0.00e+00 | 1.14e+00 | -1.90e-01 | 1.82e+00 |
| $\phi_b^{dist}$ | -1.23e-01 | 0.00e+00 | -1.17e+00 | 5.20e-01 | -1.84e+00 | 1.39e+00 |
| $\phi_b^{\Delta age}$ | -1.52e-02 | 0.00e+00 | -9.55e-01 | 6.55e-01 | -1.60e+00 | 1.60e+00 |
| $\sigma_w^{high}$ | 8.96e-01 | 9.00e-01 | 0.00e+00 | 1.76e+00 | -7.63e-02 | 2.53e+00 |
| $\sigma_w^{grow}$ | -2.87e-01 | 0.00e+00 | -1.10e+00 | 0.00e+00 | -1.67e+00 | 4.73e-01 |
| $\sigma_e^{high}$ | -4.57e-02 | 0.00e+00 | -8.86e-01 | 6.09e-01 | -1.52e+00 | 1.43e+00 |
| $\sigma_e^{grow}$ | 4.08e-02 | 0.00e+00 | -7.80e-01 | 7.01e-01 | -1.30e+00 | 1.68e+00 |
| $\sigma_d^{dist}$ | -2.00e+00 | -2.03e+00 | -3.04e+00 | -1.06e+00 | -3.59e+00 | -4.41e-01 |
| $\sigma_d^{\Delta age}$ | -1.92e-01 | 0.00e+00 | -1.05e+00 | 3.10e-01 | -1.66e+00 | 1.03e+00 |
| $\sigma_b^{dist}$ | 9.40e-02 | 0.00e+00 | -6.98e-01 | 1.15e+00 | -1.60e+00 | 1.80e+00 |
| $\sigma_b^{\Delta age}$ | 7.37e-02 | 0.00e+00 | -9.25e-01 | 8.88e-01 | -1.53e+00 | 1.94e+00 |

Table S11: **Bayes factor support for biogeographic hypotheses H1-H4**. The hypothesis column indicates associations between the four biogeographic hypotheses (main text) and the signs of feature effect parameters. Columns 3-5 report the posterior probability that a parameter was zero, positive, or negative. Column 6 gives the posterior probability that a parameter was positive (vs. negative) among non-zero-valued samples. Column 7 reports Bayes factors (BFs) supporting the feature effect had the correct signed value (+/− from column 2) or, if not associated with any hypothesis, that it had a non-zero value. Column 8 gives false discovery rates (FDRs) corresponding to Substantial, Strong, and Very Strong significance levels from Table S6. Column 9 provides the Bayes factor interpretations for model selection as suggested by Kass & Rafferty [9]: $< 1.0$ = Opposing, $\approx 1.0$ = None, $> 1.0$ = Marginal, $>3.2$ = Substantial; $> 10$ = Strong, $>32$ = Very strong. Hypothesis in bold indicate significant support (Substantial or better) from the parameter estimate on that row. Each of the four hypothesis received significant support from at least one model parameter, and no parameters generated significant opposing support. Table S6 suggests false discovery rates for all significant results are acceptably low.

| Variable | Hypothesis | $=0$ | $>0$ | $<0$ | $\frac{>0}{>0+<0}$ | BF | FDR | Interpretation |
|---|---|---|---|---|---|---|---|---|
| $\phi_w^{size}$ | H1+ | 0.54 | 0.34 | 0.12 | 0.75 | 1.5 | – | Marginal |
| $\phi_w^{age}$ | – | 0.61 | 0.30 | 0.09 | 0.76 | 2.3 | – | Marginal |
| $\phi_e^{size}$ | **H1−** | 0.30 | 0.13 | 0.57 | 0.18 | 4.0 | 4.3 | Substantial |
| $\phi_e^{age}$ | – | 0.46 | 0.35 | 0.19 | 0.65 | 1.9 | – | Marginal |
| $\phi_d^{dist}$ | **H3−** | 0.02 | 0.01 | 0.97 | 0.01 | 97.0 | 5 | Very strong |
| $\phi_d^{\Delta age}$ | **H2+** | 0.41 | 0.55 | 0.04 | 0.94 | 3.7 | 26.1 | Substantial |
| $\phi_b^{dist}$ | – | 0.48 | 0.20 | 0.32 | 0.38 | 1.1 | – | None |
| $\phi_b^{\Delta age}$ | – | 0.51 | 0.24 | 0.25 | 0.49 | 1.0 | – | None |
| $\sigma_w^{high}$ | **H4+** | 0.25 | 0.71 | 0.04 | 0.95 | 7.3 | 0 | Substantial |
| $\sigma_w^{grow}$ | H4− | 0.48 | 0.10 | 0.42 | 0.19 | 2.2 | – | Marginal |
| $\sigma_e^{high}$ | H4− | 0.53 | 0.22 | 0.25 | 0.48 | 1.0 | – | None |
| $\sigma_e^{grow}$ | H4+ | 0.52 | 0.26 | 0.22 | 0.55 | 1.1 | – | None |
| $\sigma_d^{dist}$ | **H3-** | 0.02 | 0.00 | 0.98 | 0.00 | 147.0 | 0 | Very strong |
| $\sigma_d^{\Delta age}$ | – | 0.53 | 0.15 | 0.32 | 0.32 | 0.5 | – | Marginal (opposing) |
| $\sigma_b^{dist}$ | – | 0.47 | 0.31 | 0.22 | 0.58 | 1.1 | – | None |
| $\sigma_b^{\Delta age}$ | – | 0.48 | 0.29 | 0.23 | 0.55 | 1.1 | – | None |

Table S12: **Regional net diversification rates through time.** Posterior medians and 80% highest posterior density intervals are shown. Blank entries correspond to times when that region did not exist.

| Region | Epoch 7 | Epoch 6 | Epoch 5 | Epoch 4 | Epoch 3 | Epoch 2 | Epoch 1 |
|---|---|---|---|---|---|---|---|
| Z | 0.148 [0.019, 0.288] | 0.148 [0.019, 0.288] | 0.148 [0.019, 0.288] | 0.148 [0.019, 0.288] | 0.148 [0.019, 0.288] | 0.148 [0.019, 0.288] | 0.148 [0.019, 0.288] |
| G | | 0.241 [0.052, 0.445] | 0.278 [-0.096, 0.777] | 0.162 [-0.477, 1.002] | -0.076 [-1.186, 0.597] | -0.194 [-3.054, 0.754] | -0.656 [-31.250, 2.237] |
| N | | | 0.172 [0.001, 0.332] | 0.179 [-0.161, 0.685] | -0.019 [-0.364,0.304] | -0.078 [-0.952,0.333] | -0.132 [-1.687,0.515] |
| K | | | | 0.196 [-0.033, 0.412] | 0.239 [0.026, 0.420] | 0.350 [0.077, 0.668] | 0.334 [0.069, 0.671] |
| O | | | | | 0.208 [-0.027,0.460] | 0.248 [-0.005,0.455] | 0.327 [0.077, 0.637] |
| M | | | | | | 0.229 [-0.026,0.550] | 0.325 [0.042,0.616] |
| H | | | | | | | 0.241 [-0.010,0.667] |
